# When spike sorting matters: single-neuron identity in the quantification of neural circuits

**DOI:** 10.64898/2026.11.19.767959

**Authors:** Daniel Suárez-Barrera, Norberto Encinas-Rodríguez, Héctor Díaz, Antonio Zainos, Lucas Bayones, Román Rossi-Pool

## Abstract

Multiunit threshold crossings support high-performance brain–computer interfaces and preserve the low-dimensional trajectories that summarize population dynamics, raising the question of when spike sorting is still necessary. Using previously published recordings from four rhesus monkeys, we analyzed areas 3b and 1 of primary somatosensory cortex and ventral premotor cortex during a bimodal detection task, and ventral premotor cortex during an interval comparison task. From the same spikes, we built three representations: putative single units obtained with nonlinear sorting (UMAP and HDBSCAN), intermingled units obtained with linear sorting, and unsorted multiunit activity. Nonlinear sorting resolved up to three times more units than multiunit activity. In area 1, mixing discarded the periodicity information carried by phase-locked neurons, lengthened intrinsic timescales by about 40%, increased trial-to-trial variability, and raised neurometric detection thresholds by 33%. In premotor cortex, mixing reduced the task-related information recovered from the population and hid significant dimensions of its dynamics. After geometric alignment, the overall shape of population trajectories was nearly identical across representations, but distances between population states were distorted, and the approximately linear representation of interval duration saturated for longer intervals. Multiunit activity is therefore sufficient for analyses that rely on the dominant, shared structure of population activity, such as decoding, whereas analyses of single-neuron coding, variability, timescales, dimensionality, and parametric representations often require spike sorting.

## Introduction

A central goal of systems neuroscience is to quantify how neural circuits encode sensory variables, hold information over time, and generate decisions. Extracellular electrophysiology remains central to that goal, resolving spiking activity at millisecond precision. Such analyses have long relied on spike sorting to assign extracellular waveforms to putative single neurons, preserving the single-neuron identity needed to estimate selectivity, trial-to-trial variability, noise correlations, and spike–field coupling. As intracortical recordings have grown in channel count and moved toward real-time use, that necessity has come into question.

Threshold-based approaches make a strong practical case for skipping the step. In human intracortical brain–computer interfaces, threshold crossings and related spike-band features support robust real-time decoding of intended movement, cursor control, handwriting, and speech, including high-performance brain-to-text communication in participants with paralysis (Gilja et al., 2015; Pandarinath et al., 2017; Willett et al., 2021, 2023). Population analyses point the same way: multiunit threshold crossings preserve the geometry of low-dimensional neural trajectories and yield conclusions comparable to those drawn from sorted single units (Trautmann et al., 2019; Rossi-Pool and Romo, 2019). Random projection arguments explain why. When the relevant activity lies on a smooth, low-dimensional manifold, random mixtures of neuronal activity retain the global structure of the population state space. For real-time decoding and for estimating global population dynamics, then, single-neuron identity may be dispensable.

Preserving population geometry is not the same as preserving neural coding. The critical question is not whether unsorted activity can support decoding or manifold estimation, but whether it can support the biological inference being made, and many measurements used to quantify circuit function depend on single-neuron identity. Classical neurometric analyses compare the sensitivity of individual neurons with behavioral performance to ask whether single units carry enough information to account for perception or decision making (Newsome et al., 1989; Britten et al., 1992; Hernández et al., 2000; Romo & Salinas, 2003; de Lafuente & Romo, 2005). Analyses of stimulus selectivity, choice-related activity, and mixed selectivity ask not only whether task conditions separate in population space, but which neurons carry information and how their activity predicts behavior. Correlation-based measures are defined between neuronal pairs: noise correlations depend on tuning similarity, cortical distance, and behavioral state, and they shape both the information encoded by a population and its readout by downstream circuits (Cohen & Kohn, 2011; Ecker et al., 2014; Kohn et al., 2016; Valente et al., 2021; Panzeri et al., 2022; Papale et al., 2025). Oscillatory coupling raises the same difficulty, since phase locking and rhythm-specific routing require knowing which spikes align to which local field potential phases, and which populations take part in a given interaction (Fries et al., 2001; Womelsdorf et al., 2007; Buffalo et al., 2011; Bastos et al., 2020). Pooling neurons into one multiunit signal can therefore preserve the global population state while distorting single-neuron selectivity, neurometric thresholds, pairwise correlation structure, and spike–field coupling.

Dimensionality is one population measure where this should show. Dimensionality reduction is widely used to estimate how many latent modes describe a population’s dynamics (Cunningham & Yu, 2014; Gao & Ganguli, 2015; Vyas et al., 2020), and the estimates are read as constraints on the computations a circuit implements. Low-dimensional activity suggests that heterogeneous single-neuron responses are organized by a smaller set of coordinated modes; higher-dimensional activity may reflect richer mixed selectivity or more distributed interactions. These estimates depend on how activity is sampled. Pooling spikes across neurons merges partially independent response components, shifts the balance between private and shared variance, and suppresses weak but informative patterns. Multiunit activity can preserve the global structure of population trajectories and still alter the estimated dimensionality of the circuit.

We expect this cost to scale with the heterogeneity of the neurons a multiunit signal pools, and therefore with hierarchical level. Early sensory areas contain neurons with stronger stimulus-locked responses and shorter latencies, whereas higher-order areas integrate sensory evidence with context, memory, and behavioral demands (Felleman & Van Essen, 1991; Murray et al., 2014; Rossi-Pool et al., 2021; Parra et al., 2022; Bayones et al., 2024). In primary somatosensory cortex, where neighboring neurons share receptive fields and stimulus phase preferences, multiunit activity should preserve stimulus-driven population structure while still distorting neurometric estimates and fine tuning, with the cost growing from area 3b to area 1 (Parra et al., 2022). In ventral premotor cortex (PMv), where neurons show mixed selectivity and longer integration timescales, losing single-neuron identity should have broader consequences for dimensionality, decoding, and circuit interactions (Díaz et al., 2025; Andrade-Ortega et al., 2025). Recording from two subdivisions in each region (areas 3b and 1 in S1; F4 and F5 in PMv) also lets us compare the effect of sorting within regions as well as between them.

Here we ask when spike sorting matters. We analyzed previously published recordings from four *Macaca mulatta* performing two tasks: a bimodal detection task (BDT), recorded in areas 3b and 1 of primary somatosensory cortex (Parra et al., 2022) and in PMv (Andrade-Ortega et al., 2025), and a time interval comparison task (TICT), recorded in PMv (Díaz et al., 2025). For each dataset we compared three representations of the same spikes: putative single-unit activity (PSA), obtained with a nonlinear UMAP-based method that recovers individual neurons with high precision (Suárez-Barrera et al., 2025); intermingled activity (IA), obtained with a linear sorting that partially mixes neighboring neurons; and multiunit activity (MUA), unsorted threshold crossings that pool nearby neurons without resolving them. Across these representations we quantified neural coding, variability, neurometric sensitivity, dimensionality, and population dynamics. Nonlinear sorting resolved many more neurons than either linear sorting or multiunit pooling. In area 1, single neurons were less variable, had shorter intrinsic timescales, and supported lower neurometric thresholds, and pooling discarded the periodicity information carried by their phase-locked responses. In PMv, the three representations encoded task variables differently and yielded different numbers of significant dimensions. At the population level, the geometry of neural dynamics also changed with the degree of neuronal isolation, and mixing distorted the linear representation of interval duration. These results mark the boundary between analyses that can rely on multiunit activity and those that require single-neuron identity.

## Results

### Three representations of the same extracellular spikes

A microelectrode records action potentials from several neurons near its tip, and these spikes differ in amplitude and waveform shape (Fig. 1A). We compared three representations of this activity, which differ in how much of that waveform information they use (Fig. 1). Spike sorting uses those differences to assign each spike to a putative single neuron. Alternatively, the recording can be analyzed without sorting by detecting threshold crossings of the band-pass-filtered signal, which provide a measure of multiunit activity (MUA; see Methods). MUA combines the activity of nearby neurons without resolving their individual identities, and it has been used to relate spiking to local field potentials (Bastos et al., 2020), to characterize population dynamics (Trautmann et al., 2019), and to drive brain–computer interfaces (Gilja et al., 2015; Willett et al., 2021, 2023).

**Figure 1.**
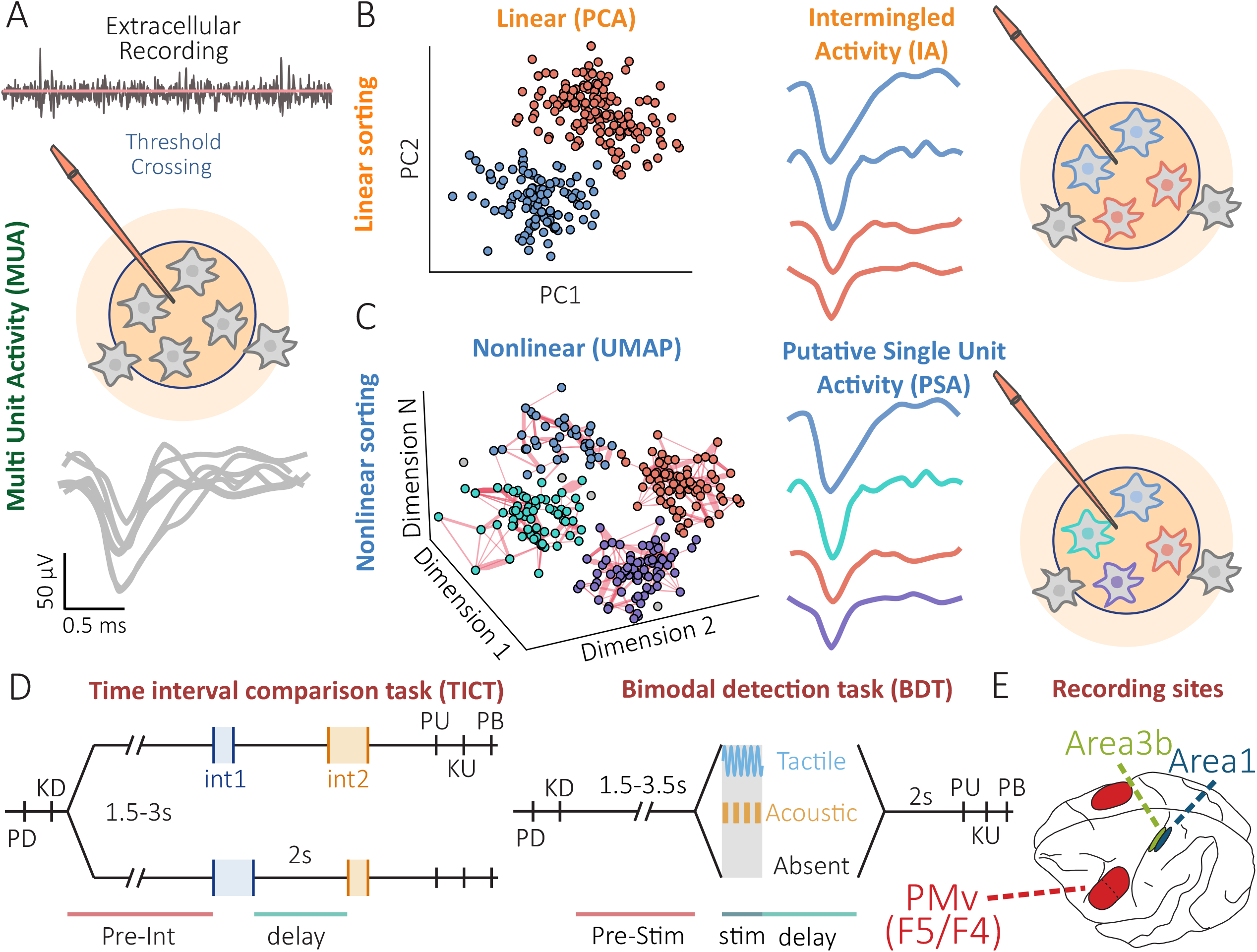
Spike sorting approaches, behavioral tasks, and cortical recording sites. (A) Schematic of an extracellular recording obtained with a microelectrode. Action potentials from several nearby neurons contribute to the recorded signal. Threshold crossings quantify multiunit activity (MUA) without identifying individual neurons. The waveforms below illustrate spikes from different neurons. (B) Linear spike sorting based on principal component analysis (PCA). Spike waveforms are projected onto the first two principal components (PC1 and PC2), where two clusters are identified. Spikes from neurons with similar waveforms can fall within the same cluster, producing intermingled activity (IA). The waveforms and schematic on the right show spikes from different neurons grouped within the same units. (C) Nonlinear spike sorting using UMAP for dimensionality reduction followed by HDBSCAN for clustering (Suárez-Barrera et al., 2025). Applied to the same spikes shown in B, this approach identifies four clusters instead of two. Because the identity of individual neurons cannot be confirmed from extracellular recordings alone, we refer to this representation as putative single-unit activity (PSA). (D) Schematic of the two behavioral tasks. Left: time interval comparison task (TICT), in which monkeys compared the durations of two tactile intervals (int1, blue; int2, orange) separated by a fixed 2-s delay. After the second interval and a subsequent delay, the animals indicated which interval was longer by pressing one of two buttons (Díaz et al., 2025). Right: bimodal detection task (BDT), in which monkeys reported whether a tactile stimulus (blue), an acoustic stimulus (orange), or no stimulus (gray) had been presented. Stimuli lasted 500 ms and were followed by a 2-s delay before the behavioral response (Parra et al., 2022; Andrade-Ortega et al., 2025). PD, probe down; KD, key down; PU, probe up; KU, key up; PB, push button. (E) Schematic of the recording locations in the macaque brain: areas 3b and 1 of primary somatosensory cortex, and ventral premotor cortex (PMv), including its caudal (F4) and rostral (F5) parts; PMv was recorded in both hemispheres (red ovals).

A common approach to spike sorting uses linear dimensionality reduction, such as principal component analysis (PCA), to separate neurons by their spike waveforms; here, the PCA embedding was clustered with HDBSCAN, as for nonlinear sorting (see Methods). As we showed previously (Suárez-Barrera et al., 2025), these methods identify distinct units but often fail to separate neurons whose waveforms are similar. In Fig. 1B, PCA yields two clusters, each merging the spikes of two neurons with similar waveforms (blue and red). We call this mixture of spikes from different neurons within a single unit intermingled activity (IA). Linear sorting thus resolves more units than the single multiunit signal on each channel, yet some of those units still combine several neurons.

Nonlinear dimensionality reduction separates spikes from different neurons more effectively. On the same spike waveforms, we applied UMAP for dimensionality reduction followed by HDBSCAN for clustering (Fig. 1C; see Methods). We previously showed, with synthetic datasets and paired intracellular and extracellular recordings, that this approach identifies individual neurons more accurately than linear methods and is less likely to merge spikes from different neurons into a single unit (Suárez-Barrera et al., 2025). In the example shown, nonlinear sorting identifies four putative neurons (cyan, red, purple, and blue), whereas PCA-based sorting finds only two units. We call the resulting activity putative single-unit activity (PSA), since some units may still contain spikes from more than one neuron. Our aim is not to benchmark sorting algorithms but to determine how the degree of neuronal isolation shapes the characterization of cortical circuits during cognitive tasks. We therefore compare three representations of extracellular activity: multiunit activity (MUA), intermingled activity (IA), and putative single-unit activity (PSA). All three derive from the same spikes on each channel and differ only in how those spikes are grouped: MUA contains every spike that sorting distributes among the units of a channel, plus those it leaves unassigned.

### Behavioral tasks and cortical recording areas

To examine how spike sorting affects the characterization of neural circuits, we reanalyzed extracellular recordings from two previously studied perceptual tasks. In the first, a bimodal detection task (BDT), two rhesus monkeys (*Macaca mulatta*) were trained to detect tactile or acoustic stimuli presented at different intensities, including near-threshold levels (Parra et al., 2022; Andrade-Ortega et al., 2025). Each trial began with a variable foreperiod of 1.5–3.5 s, followed by a 500-ms tactile stimulus, an acoustic stimulus, or no stimulus (Fig. 1D, right). Tactile stimuli were 20-Hz vibrations of varying amplitude; acoustic stimuli were 1-kHz tones amplitude-modulated at 20 Hz and presented at different intensities. After the stimulus period, the monkeys held their decision through a 2-s delay and then reported, by pressing one of three buttons, whether they had perceived a tactile stimulus, an acoustic stimulus, or none. Because stimulus presentation and the behavioral response are separated in time, the task lets us examine both sensory coding and the maintenance of perceptual decisions. In figures showing BDT conditions, tactile and acoustic stimuli appear in blue and orange, respectively.

Previous work with this task revealed differences in neural processing across cortical areas. In the primary somatosensory cortex (S1), Parra et al. (2022) showed that neurons in areas 3b and 1 encoded tactile information but differed in response latency, periodicity information, and neuronal variability, differences that pointed to a functional hierarchy within S1. In the ventral premotor cortex (PMv), Andrade-Ortega et al. (2025) found neurons that encoded both sensory modalities, along with activity that maintained the encoding of the bimodal decision throughout the delay. At the population level, sensory and maintenance-related activity followed distinct dynamics. These findings make it possible to ask whether spike sorting has different consequences in circuits involved in different stages of sensory processing and decision making.

The recordings from these studies are publicly available on Zenodo (Zainos et al., 2022, 2024a). The dataset included 54 electrode recordings from area 3b, 77 from area 1, and 531 from PMv (Fig. 1E). To examine possible differences within PMv, we further divided these recordings into 382 from its rostral part (F5) and 149 from its caudal part (F4), following the nomenclature of Matelli et al. (1985). These recordings let us compare the effects of spike sorting across sensory and premotor areas, and between subdivisions of the same region.

We also analyzed recordings from a time interval comparison task (TICT), in which two rhesus monkeys were trained to compare the durations of two consecutive intervals (Díaz et al., 2025; Suárez-Barrera et al., 2025b; Encinas-Rodríguez et al., 2025). Each trial began with a variable foreperiod of 1.5–3 s, followed by a first interval (int1) bounded by two brief tactile pulses (Fig. 1D, left). The duration of int1 ranged from 400 to 2,000 ms. After a fixed 2-s delay, a second interval (int2) was presented, and after a further delay the monkeys indicated which interval was longer (int1 > int2 or int2 > int1) by pressing one of two buttons. Unlike the BDT, where the animals held a perceptual decision through the delay, the TICT required them to retain the duration of the first interval for comparison with the second.

Díaz et al. (2025) previously analyzed these recordings and showed that neurons in PMv encoded interval duration through heterogeneous responses. At the population level, a parametric, approximately linear representation of duration emerged toward the end of int1 and persisted throughout the 2-s delay, so that the duration of the first interval could be read out from the population state. Here we analyzed 311 electrode recordings from PMv during this task, publicly available on Zenodo (Zainos et al., 2024b). These data let us examine how spike sorting affects the characterization of temporal coding and population dynamics in a premotor circuit engaged in interval estimation and working memory.

### Spike sorting changes firing rates, unit counts, and population composition

We first examined how each sorting approach affected firing-rate distributions and the number of identified units across cortical areas. Figure 2 shows the cumulative distributions of firing rates for PSA, IA, and MUA. In every area and in both tasks, PSA units had lower firing rates than IA and MUA units (Fig. 2A–E, left), consistent with our previous finding that nonlinear sorting recovers low-firing neurons whose spikes would otherwise stay mixed with those of other neurons (Suárez-Barrera et al., 2025).

**Figure 2.**
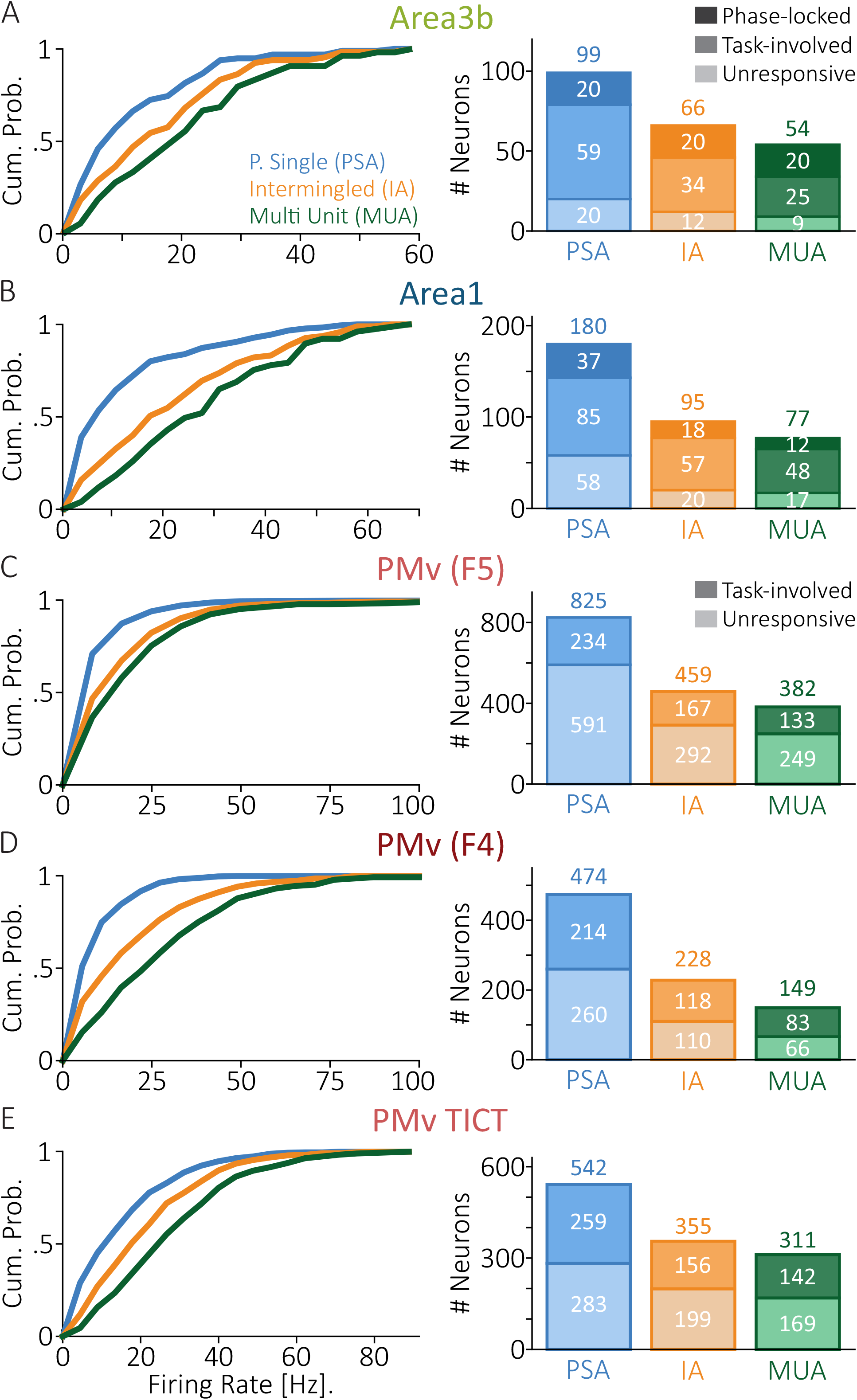
Effects of spike sorting on firing rates and neuronal populations across cortical areas. (A–E) Firing-rate distributions and numbers of identified units for three representations of extracellular activity: putative single-unit activity (PSA, blue), intermingled activity (IA, orange), and multiunit activity (MUA, green). Left panels show the cumulative distributions of firing rates for each representation. Right panels show the total number of identified units and their functional classification. Numbers above the bars give the total number of units; numbers within each segment give the number of units in each category. (A, B) Recordings from areas 3b (A) and 1 (B) of primary somatosensory cortex during the bimodal detection task (BDT). Units were classified as phase-locked to the periodic tactile stimulation, task-involved, or unresponsive, shown as different shades within each bar. (C, D) Recordings from PMv (F5) (C) and PMv (F4) (D) during the BDT. Units were classified as task-involved or unresponsive. (E) Recordings from PMv during the time interval comparison task (TICT), with units classified as in C and D.

The three approaches also differed markedly in how many units they identified (Fig. 2A–E, right). PSA always yielded the most units, IA intermediate numbers, and MUA the fewest. In somatosensory cortex, the count rose from 54 MUA channels to 99 PSA units in area 3b, and from 77 to 180 in area 1. PMv followed the same pattern during the BDT, with PSA identifying 825 units in F5 and 474 in F4, against 382 and 149 MUA channels. During the TICT, PSA yielded 542 units in PMv from 311 MUA channels. Overall, PSA increased the number of units 1.7- to 3.2-fold relative to MUA, with the largest increase in F4.

These differences extended to the functional properties of the identified populations. In somatosensory cortex, units were classified as task-involved, phase-locked to the periodic stimulation, or unresponsive (see Methods). PSA identified more task-involved units than IA and MUA in both areas. Phase-locked units also increased in area 1, from 12 with MUA to 37 with PSA, but in area 3b their number stayed at 20 across the three approaches, so their proportion fell from 37% to 20% as additional, non-phase-locked units were resolved. In PMv, PSA likewise identified more task-involved units in both the BDT and the TICT. The increase was not limited to task-related activity, however: PSA also yielded more unresponsive units. Spike sorting therefore changes the number and firing-rate distribution of the identified units, as well as the composition of the recorded populations. The following sections examine whether these differences affect estimates of neural coding and population dynamics.

### Spike sorting reveals neuronal coding masked by mixed activity

We then examined how spike sorting affects the coding properties of individual neurons. Figure S1A–C shows several examples from PMv during the BDT in which PSA identified neurons with distinct response profiles that IA and MUA did not resolve. In these recordings, IA and MUA merged neurons with different sensory preferences and temporal responses into fewer units with higher firing rates, consistent with the heterogeneity of single-unit responses in premotor cortex, where individual neurons mix temporal and categorical dynamics (Diaz-deLeon et al., 2022). The same happened during the TICT (Fig. S1D), where neurons with distinct temporal profiles contributed to the same mixed activity.

Figure 3A shows an example from area 1 in which a putative single neuron responded strongly to tactile stimulation, with a relatively low baseline firing rate and clear stimulus-locked activity. The corresponding IA and MUA representations showed higher firing rates. Despite its lower firing rate, the neuron identified with PSA carried more information about stimulus periodicity, as measured by mutual information (MI; see Methods). This information was markedly reduced in IA and MUA, suggesting that mixing spikes from different neurons can obscure the coding of specific stimulus features. Area 3b showed a similar example (Fig. S2A), with the three representations again differing in firing rate and periodicity information.

**Figure 3.**
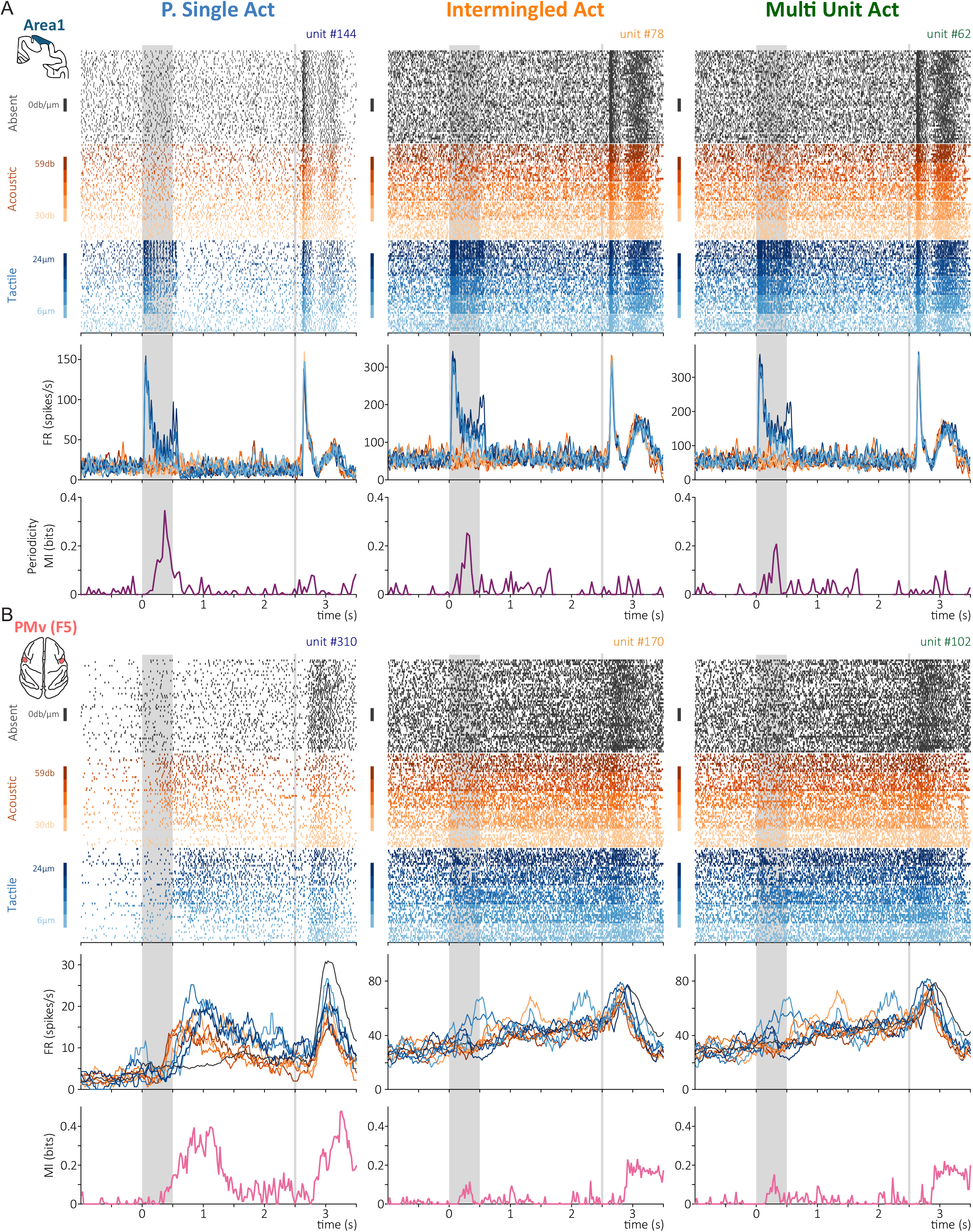
Spike sorting reveals differences in sensory coding and information content. (A) Area 1 during the BDT. The same recording analyzed as PSA (left), IA (middle), and MUA (right). From top to bottom: spike rasters, firing rates, and MI about stimulus periodicity. Trials are grouped by condition: absent (black), acoustic (orange; 30 and 59 dB), and tactile (blue; 6 and 24 μm), with lighter shades for lower intensities. The PSA neuron had a lower firing rate and more periodicity information than the IA and MUA units. (B) PMv (F5) during the BDT, organized as in A, with MI about stimulus modality in the bottom panels. The PSA neuron distinguished tactile and acoustic stimulation from stimulus absence, mainly during the early delay; this information disappeared in IA and MUA despite their higher firing rates. In A and B, gray shading marks the stimulus period, vertical gray lines mark the response cue, and time is relative to stimulus onset.

Differences were also apparent in PMv. Figure 3B shows an example from F5 in which PSA identified a neuron that clearly distinguished trials with tactile or acoustic stimulation from trials without a stimulus, and carried information about stimulus modality mainly during the first part of the delay. In IA and MUA, firing rates were higher, but this information disappeared: the activity of additional neurons diluted the stimulus-related information carried by the isolated neuron.

A similar effect appeared in the persistent activity of F4 (Fig. S2B). The neuron identified with PSA maintained a clear difference between stimulus conditions throughout the delay, carrying sustained modality information driven mainly by acoustic trials. In IA, this information was reduced. In MUA, the signal also included activity from neurons responding near the response period, which changed the temporal profile of information. As a result, the persistent coding seen with PSA became less apparent in the mixed representations.

These examples show that higher firing rates do not necessarily mean more information about task variables. Mixing spikes from different neurons can reduce apparent selectivity, alter the time course of information, and obscure responses that spike sorting resolves. We next examined whether these differences extend across the recorded populations.

### Spike sorting differentially affects neuronal coding across the somatosensory hierarchy

Previous studies have revealed a functional hierarchy between areas 3b and 1 of the primary somatosensory cortex. Neurons in area 3b carry more information about the periodic structure of vibrotactile stimuli and have shorter response and coding latencies than those in area 1, whereas firing-rate information does not differ between the two areas (Parra et al., 2022). Area 1 neurons also have longer intrinsic timescales and greater trial-to-trial variability (Parra et al., 2022; Tauste Campo et al., 2024). During tactile stimulation, area 1 shows an increase in intra-area functional interactions that is absent in area 3b, and neurons with longer timescales take part in more of these interactions, pointing to a greater role for local recurrent processing in area 1 (Tauste Campo et al., 2024). The two areas thus differ in temporal integration and functional interactions as well as in sensory coding. We therefore hypothesized that coding in area 1 would be more sensitive to spike sorting than in area 3b: if neighboring neurons in area 1 are more heterogeneous, pooling them should degrade their coding more.

We first examined how spike sorting affected the coding of stimulus periodicity. In area 1, PSA identified more neurons carrying significant periodicity information throughout the stimulation period than IA or MUA (Fig. 4A). Summed periodicity information was also higher with PSA, whereas IA and MUA reached similar, lower values (Fig. 4B). This difference was not just a consequence of the larger number of units: the cumulative distributions show that individual PSA neurons also carried more periodicity information than IA or MUA units (Fig. 4E, left). In area 3b, sorting had a weaker effect. PSA again identified more neurons carrying periodicity information, but the differences in summed information were small (Fig. S3A,B), and the cumulative distributions largely overlapped across the three representations (Fig. 4E, right). Periodic coding in area 3b was therefore robust to spike mixing, whereas mixing reduced the periodicity information recovered from area 1.

**Figure 4.**
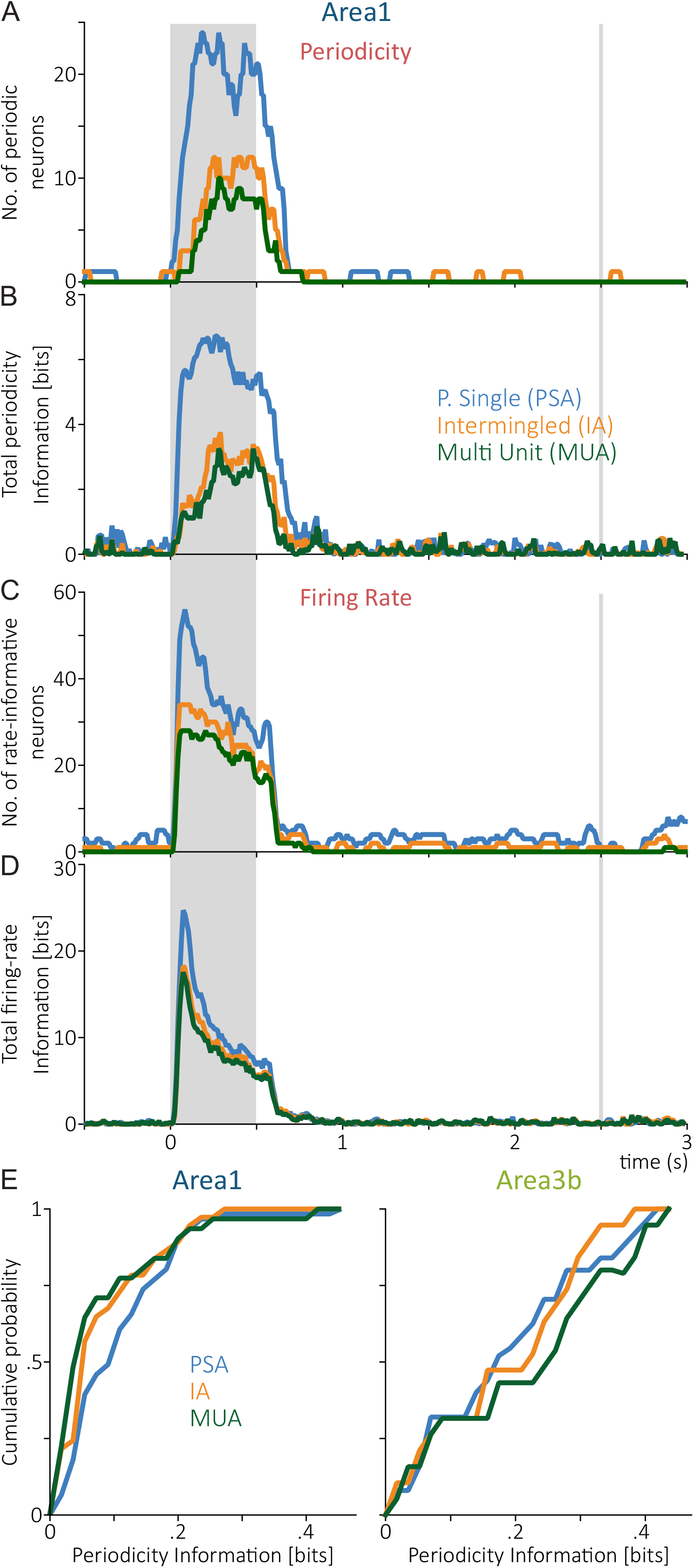
Effects of spike sorting on periodicity and firing-rate coding. (A–D) Area 1 during the BDT, for PSA (blue), IA (orange), and MUA (green). (A) Number of units with significant periodicity information over time. (B) Total periodicity information, summed across units. (C) Number of units with significant firing-rate information about stimulus amplitude. (D) Total firing-rate information, summed across units. (E) Cumulative distributions of periodicity information per unit in area 1 (left) and area 3b (right). In A–D, time is relative to stimulus onset; gray shading marks the 500-ms stimulus period, and vertical gray lines mark the response cue after the 2-s delay.

We next examined whether these differences extended to firing-rate coding. In area 1, PSA identified more neurons carrying significant firing-rate information than IA or MUA, particularly during stimulus presentation (Fig. 4C). Summed firing-rate information was also higher with PSA, mainly at stimulus onset, and the differences between sorting approaches were smaller than for periodicity information (Fig. 4D). Area 3b showed a similar but weaker effect: PSA identified more rate-informative neurons, while summed information was similar across the three representations (Fig. S3C,D). The cumulative distributions, however, revealed a pattern opposite to that of periodicity. In both areas, individual PSA units carried less firing-rate information than IA or MUA units, with a larger difference in area 1 (Fig. S3E). The higher summed information obtained with PSA therefore reflected a larger number of informative neurons rather than more information per unit. Pooling spikes thus increased the rate information carried by each unit while reducing periodicity information, so the effect of spike sorting depends on the type of information being quantified as well as on the cortical area.

Neurons recorded on the same electrode can differ in the timing of their responses to periodic stimulation. When neurons with different preferred phases are combined, their periodic modulations partially cancel, so the pooled signal loses periodicity information even if each neuron carries a great deal of it. Pooling neurons with similar firing-rate modulations, by contrast, preserves or even increases the rate information of the mixed signal. This interpretation fits the larger effect of sorting in area 1, where response latencies are longer and more dispersed than in area 3b (Parra et al., 2022), and neighboring neurons are therefore less likely to share the same phase. The impact of spike sorting therefore cannot be reduced to the number of units recovered; it depends on the temporal and functional heterogeneity of the neurons that contribute to each recording.

### Spike sorting affects neuronal variability and intrinsic timescales

Given the differences in periodicity coding observed in area 1, we next asked whether spike sorting also affected measures of neuronal variability and temporal fluctuations. We first estimated the intrinsic autocorrelation timescale (τ; see Methods), which is used to characterize temporal integration and to establish functional hierarchies across cortical circuits (Murray et al., 2014; Chaudhuri et al., 2015; Rossi-Pool et al., 2021; Parra et al., 2022; Tauste Campo et al., 2024). In area 1, the estimated timescale depended on the sorting method (Fig. 5A). PSA yielded a shorter timescale (τ = 35.7 ± 5.0 ms) than IA (44.8 ± 5.8 ms) or MUA (49.9 ± 6.0 ms; errors estimated by bootstrap), and the difference between PSA and MUA was significant (bootstrap, p < 0.01). MUA therefore produced an estimate about 40% longer than PSA. The cumulative distribution of individual timescales shifted accordingly toward shorter values with PSA (Fig. 5A, right; Kolmogorov–Smirnov (KS) test with bootstrap resampling, p = 0.0328). Spike mixing can thus inflate estimates of intrinsic neuronal timescales, and differences of this size could change conclusions about temporal integration and cortical hierarchies, particularly when comparing brain areas or studies that rely on different spike sorting procedures.

**Figure 5.**
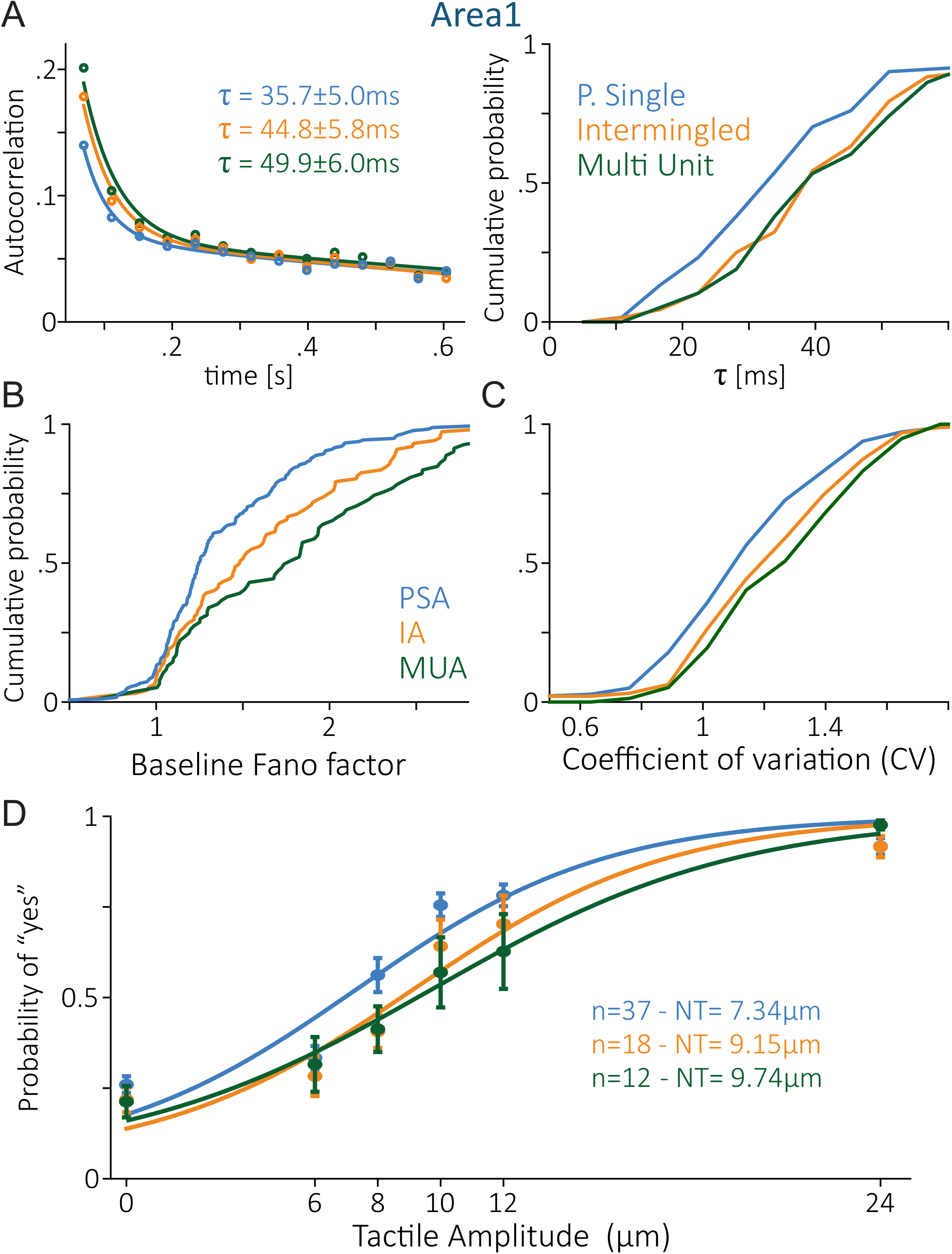
Spike sorting affects neuronal timescales, variability, and neurometric sensitivity. All panels show area 1 during the BDT. (A) Left, average autocorrelation functions for PSA (blue), IA (orange), and MUA (green), with the fitted intrinsic timescale of each representation (errors estimated by bootstrap; PSA vs. MUA, p < 0.01). Right, cumulative distributions of the timescales of individual units (PSA vs. MUA, KS, p = 0.0328). (B) Cumulative distributions of baseline Fano factors (PSA, n = 180; IA, n = 95; MUA, n = 77; PSA vs. MUA, KS, p < 0.0001). (C) Cumulative distributions of the coefficient of variation (CV) of interspike intervals (PSA vs. MUA, KS, p = 0.0019). (D) Neurometric functions computed from the response periodicity (relative power around the 20-Hz stimulation frequency, 18–22 Hz) of area 1 units with significant periodicity coding (Fig. 4), showing the probability of a “yes” response as a function of tactile stimulus amplitude. Neurometric thresholds (NT) were 7.34 ± 0.52 μm for PSA (n = 37), 9.15 ± 1.00 μm for IA (n = 18), and 9.74 ± 1.49 μm for MUA (n = 12); errors are bootstrap standard errors.

We next examined the baseline Fano factor, a measure of trial-to-trial variability defined as the variance of spike counts divided by their mean (Churchland et al., 2010; see Methods). Like the intrinsic timescale, the Fano factor increases from area 3b to area 1 and has been used to characterize functional hierarchies (Parra et al., 2022; Tauste Campo et al., 2024). In area 1, the Fano factor distribution shifted toward lower values with PSA relative to IA and MUA (Fig. 5B; KS, p < 0.0001, PSA vs. MUA), and the mean dropped from about 1.8 with MUA to 1.4 with PSA, a 22% reduction. We then examined the coefficient of variation (CV; see Methods) of interspike intervals, which captures the temporal irregularity of firing rather than trial-to-trial fluctuations in spike counts. PSA again shifted the distribution toward lower values, though less than for the Fano factor (Fig. 5C; KS, p = 0.0019, PSA vs. MUA): the mean CV fell from about 1.28 with MUA to 1.18 with PSA. Putative single neurons were thus less variable than mixed activity, whether variability was measured across trials or within spike trains. Differences of this size call for caution when interpreting neuronal variability or comparing cortical areas without accounting for spike sorting.

Sorting also changed estimates of neuronal sensitivity to tactile stimulation. Following previous neurometric approaches (de Lafuente and Romo, 2005; Parra et al., 2022), we used the periodicity of the response, measured as the relative power around the 20-Hz stimulation frequency (18–22 Hz), to discriminate stimulus-present from stimulus-absent trials (see Methods), including all area 1 units with significant periodicity coding (Fig. 4). The resulting neurometric functions differed across sorting methods (Fig. 5D): detection thresholds rose from 7.34 μm with PSA to 9.15 μm with IA and 9.74 μm with MUA, a 33% increase from PSA to MUA. Besides lengthening intrinsic timescales and inflating variability, mixing neuronal activity therefore reduced the apparent sensitivity of area 1 neurons to tactile stimulation. Spike sorting can thus shift the characterization of variability and sensory coding enough to support different conclusions about the functional properties of the same cortical circuit.

### Spike sorting alters task-related coding in premotor cortex

In PMv (F4 and F5; Matelli et al., 1985), we asked how spike sorting affects neuronal coding. In both the BDT and the TICT, we counted the units carrying significant firing-rate information (see Methods) about stimulus modality (BDT) or interval duration (TICT), and summed that information across units (Fig. 6). Following Andrade-Ortega et al. (2025), we first looked at how PMv activity represents task variables during stimulus presentation, the delay period, and motor execution. The three sorting methods differed in both the number of informative units (Fig. 6A,C) and their summed firing-rate information (Fig. 6B,D). In F5, the differences peaked during stimulus presentation, when PSA yielded more informative units and more summed information than IA or MUA. In F4 they were larger still: they persisted through the delay and peaked during motor execution, and the sustained information seen with PSA shrank once neuronal activity was mixed.

**Figure 6.**
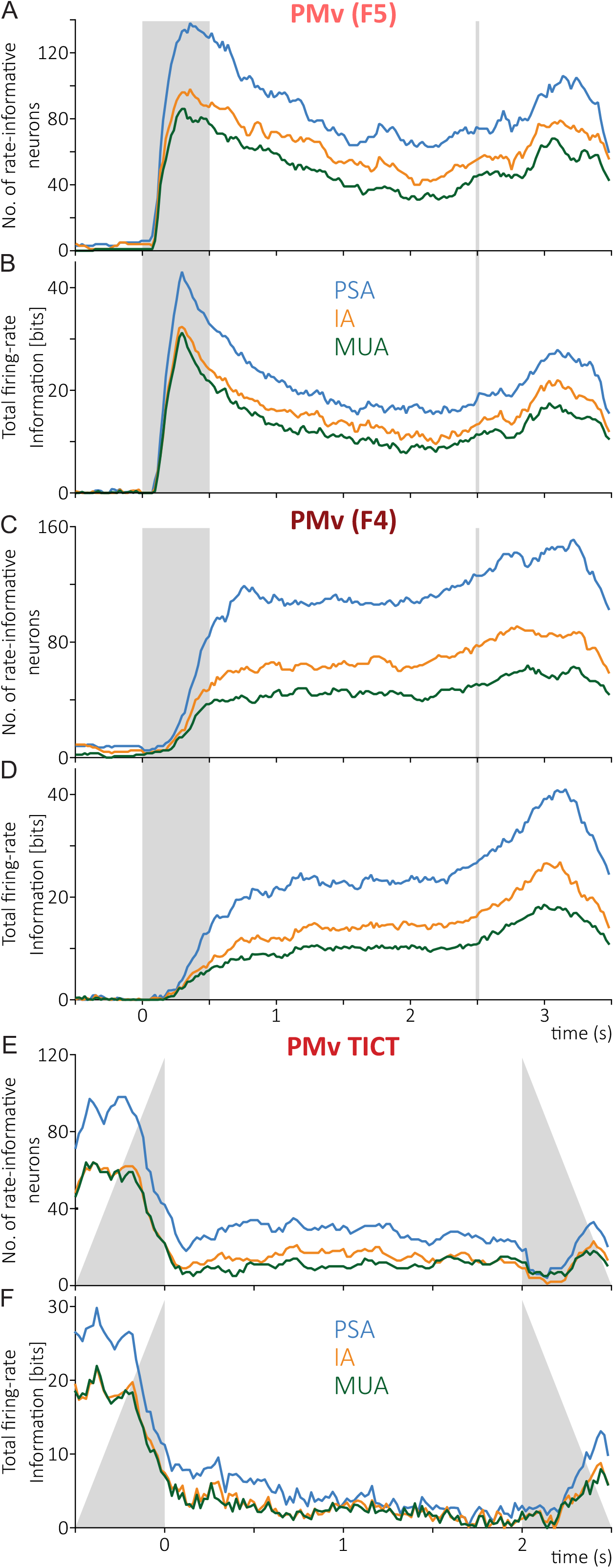
Spike sorting affects the amount and temporal distribution of task-related information in premotor cortex. (A, B) PMv (F5) during the BDT: number of units with significant firing-rate information about stimulus modality (A) and summed information across units (B). (C, D) The same measures for PMv (F4) during the BDT. (E, F) PMv during the TICT: number of units with significant firing-rate information about interval duration (E) and summed information across units (F). PSA, blue; IA, orange; MUA, green; all three were computed from the same recordings. In A–D, time is relative to stimulus onset, gray shading marks the stimulus period, and the vertical gray line marks the response cue. In E and F, time is relative to the end of int1, and gray shading marks the variable timing of int1 and int2.

The same held for interval coding during the TICT (Díaz et al., 2025). PSA again yielded more informative units and more summed information about interval duration, especially during the first interval (Fig. 6E,F), with smaller differences in the delay that followed. Because the three methods were applied to the same extracellular recordings, the extra information recovered with PSA came from resolving activity that stayed mixed in IA and MUA. Spike sorting therefore changes how much task-related information is recovered from a cortical circuit, as well as its apparent distribution and time course, and different sorting methods can lead to different conclusions about the functional organization of the same premotor circuit.

### Spike sorting alters neuronal variability and population dimensionality

We next asked whether the higher variability of mixed activity extended beyond area 1 (Fig. 5). In all three datasets (F5 and F4 during the BDT, and PMv during the TICT), the cumulative distributions shifted toward lower values with PSA relative to mixed activity (Fig. 7A; KS, p < 0.0001 in all three). The greater variability of mixed activity was therefore not restricted to sensory cortex, but also appeared in premotor circuits during both tasks.

**Figure 7.**
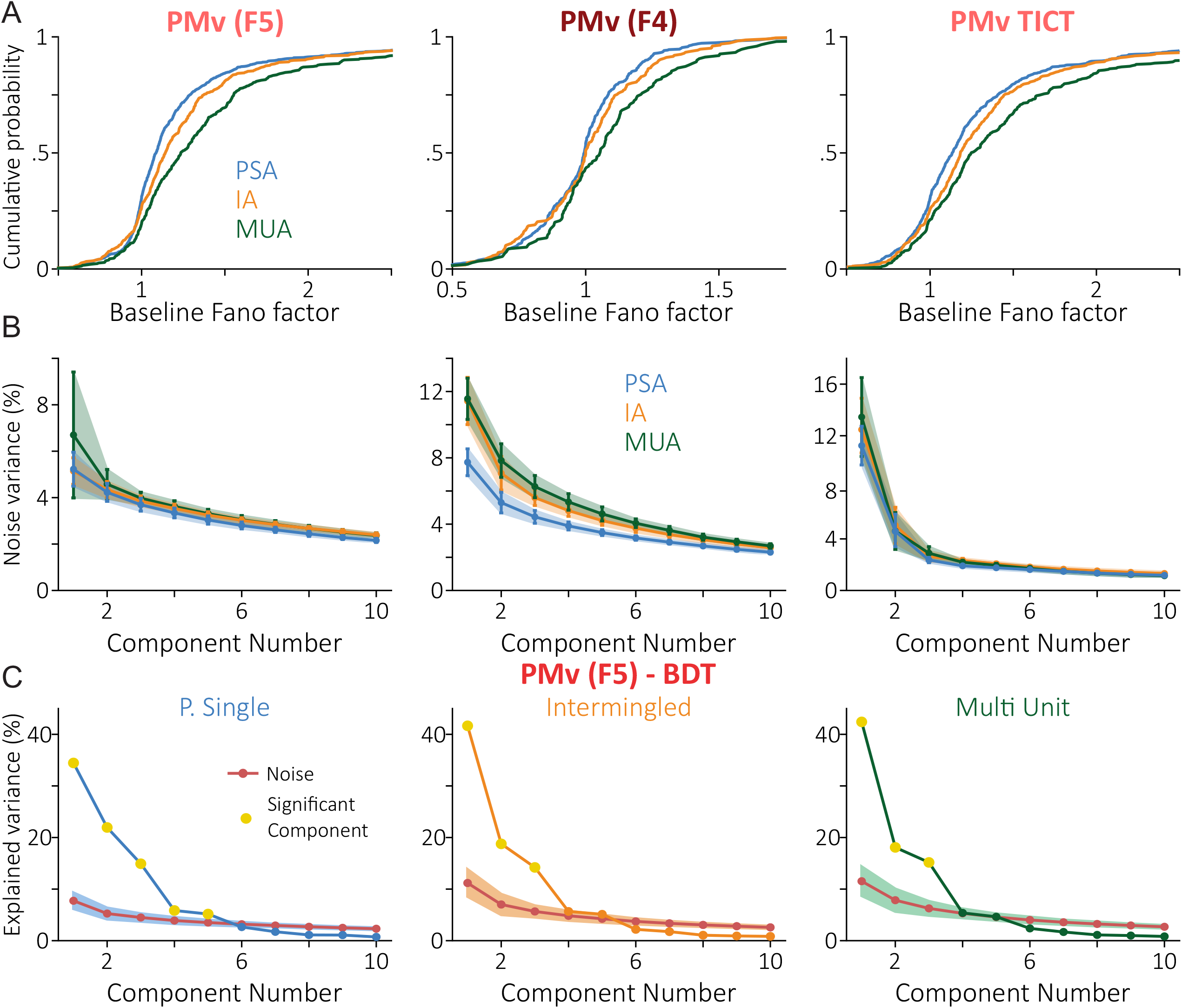
Spike sorting affects neuronal variability and the dimensionality of population dynamics in premotor cortex. (A) Cumulative distributions of baseline Fano factors in PMv (F5) and PMv (F4) during the BDT and PMv during the TICT. Sample sizes were PMv (F5) BDT (PSA, n = 825; IA, n = 459; MUA, n = 382), PMv (F4) BDT (PSA, n = 474; IA, n = 228; MUA, n = 149), and PMv TICT (PSA, n = 542; IA, n = 355; MUA, n = 311). PSA showed lower variability than MUA in all three datasets (KS, p < 0.0001 for each comparison). (B) Variance explained by the first ten noise components, estimated from trial-to-trial residuals (100 repetitions; see Methods), as a function of component number for the same three datasets. This noise variance provides the reference for identifying significant principal components. (C) PCA of PMv (F5) activity during the BDT, with responses organized by task condition and time. Explained variance is plotted against component number for each sorting method. Red curves show the variance expected from noise, and yellow markers identify significant principal components exceeding this reference. PSA revealed five significant components, compared with three for IA and MUA. PSA, blue; IA, orange; MUA, green. All analyses were performed on the same extracellular recordings processed with the three sorting methods.

This added variability could also affect the identification of significant population dimensions with principal component analysis (PCA; see Methods). PCA finds orthogonal axes that capture progressively smaller fractions of population variance, and when applied to neuronal responses organized by task condition and time, these components describe the structure of task-related population dynamics, which typically condenses into a few significant dimensions (Rossi-Pool and Romo, 2019). Their significance, however, has to be established against the variance expected from noise (see Methods; Rossi-Pool et al., 2017; Elsayed and Cunningham, 2017): we estimated the noise covariance from trial-to-trial residuals and considered a component significant only if it explained more variance than the corresponding noise component. We therefore tested whether the greater trial-to-trial variability of mixed activity came with higher noise variance, which would raise the reference that significant components must exceed. That is what we found: MUA consistently produced more noise variance than PSA, with IA generally in between (Fig. 7B), so spike sorting changed the reference against which population dimensions are judged.

To see what this means for population structure, we performed PCA on the F5 population during the BDT. Although the three methods were applied to the same extracellular recordings, PSA revealed five significant principal components, whereas IA and MUA revealed only three each (Fig. 7C). Because every population contained hundreds of units, far more than the number of task-related dimensions, insufficient sampling is unlikely to explain the difference (Gao and Ganguli, 2015). Mixing neuronal activity therefore hid two of the task-related dimensions resolved with PSA, so spike sorting shaped the dimensionality of population activity as well as the information recovered from individual units (Fig. 6). The choice of sorting procedure can thus change conclusions about the complexity and organization of cortical population dynamics, even when the underlying recordings are identical.

### Spike sorting affects the geometry of population dynamics

We finally asked whether spike sorting also affected estimates of population dynamics. PCA of the PMv population during the BDT revealed seven significant components with PSA and six with IA and MUA (Fig. S4A); the seventh carried sustained delay activity that was no longer significant with MUA. To quantify how much the population dynamics differed across sorting methods, we first computed the geometric distortion (ε95; see Methods), the smallest relative distortion of pairwise distances between population states, after optimal global scaling, that holds for at least 95% of state pairs (Trautmann et al., 2019). Because it depends only on distances, ε95 is unaffected by rotation, translation, or scaling. Both IA and MUA showed marked distortion relative to PSA (Fig. 8A, left), so distances between population states were not fully preserved. We then measured the relative squared distance (d²) between corresponding trajectories (see Methods), which was much larger for PSA–IA and PSA–MUA than for IA–MUA (Fig. 8A, middle). After Procrustes alignment, however, these distances dropped to near zero (Fig. 8A, right): most of the difference between whole trajectories could be accounted for by translating, rotating, and scaling them, even though the distances between population states remained distorted. For these comparisons we used seven components in all three representations, even though the seventh was not significant with IA or MUA. Fig. S4C,D shows examples of PCA trajectories before and after alignment, separately for the stimulus and delay periods.

**Figure 8.**
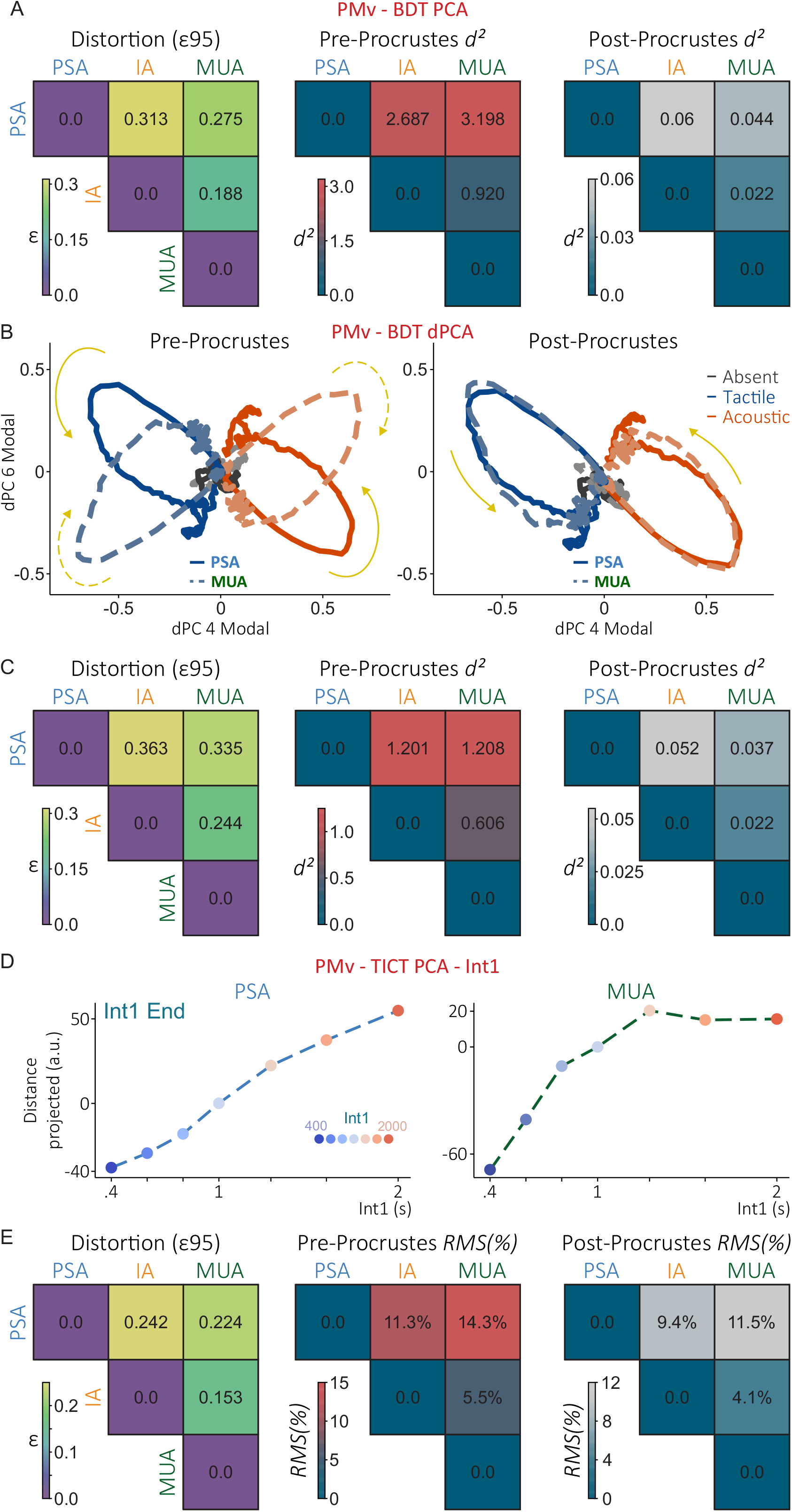
Spike sorting affects the geometry of population dynamics during sensory detection and interval comparison. (A) PCA population geometry of PMv during the BDT for PSA, IA, and MUA, computed with seven principal components. Left, geometric distortion (ε95), the smallest relative distortion of pairwise distances between population states, after optimal global scaling, that holds for at least 95% of state pairs (Trautmann et al., 2019). Middle and right, relative squared distance (d²) between corresponding trajectories before and after Procrustes alignment. (B) Population trajectories projected onto the two modal dPCA components (dPC 4 and dPC 6), before (left) and after (right) Procrustes alignment. Blue, tactile; orange, acoustic; gray, stimulus absent. Solid lines show PSA and dashed lines MUA; arrows indicate the direction of each trajectory. (C) The same comparisons as in A, computed with seven dPCA components. (D) Representation of interval duration at the end of int1 during the TICT. Projected population distance is plotted against int1 duration for PSA (left) and MUA (right); colors indicate int1 duration, from 400 to 2,000 ms. (E) Geometric distortion (ε95) and relative root-mean-square (RMS) distance between PCA representations during *int1*, before and after Procrustes alignment. In A, C, and E, the matrices show pairwise comparisons between the three representations.

We next ran a similar analysis with demixed PCA (dPCA; Kobak et al., 2016; Diaz-deLeon et al., 2022; see Methods), which separates activity shared across stimulus modalities from activity that distinguishes tactile from acoustic stimulation (Fig. S4B). As with PCA, the seventh component was not significant with MUA. Projected onto the two modal components (dPC 4 and dPC 6), PSA and MUA showed similar rotational dynamics, although before alignment their trajectories occupied different regions of population space (Fig. 8B, left). After alignment, the trajectories largely overlapped (Fig. 8B, right). ε95 and d² followed the same pattern as with PCA (Fig. 8C): geometric distortion between PSA and mixed activity, but much smaller distances between trajectories after alignment. Both PCA and dPCA therefore show that spike sorting changes the distances between population states, even though the overall shape of the dynamics is highly conserved, as the near-zero distances after alignment show.

During the TICT, we then asked how spike sorting affected population dynamics in PMv. We analyzed separately the representation of the first interval (int1) and the maintenance of its duration during the following delay (Fig. S5A,B). In both periods, PSA revealed more significant population dimensions than IA or MUA (four vs. three; Fig. S5C,D), and the fourth principal component carried task-related dynamics that were significant only with PSA (Fig. S5E,F). Several properties of the population trajectories were nonetheless similar across sorting methods. Absolute trajectory speed, as well as relative speed and relative distances estimated with KiNeT (Remington et al., 2018; see Methods), followed comparable patterns (Fig. S5G,H,J), so much of the temporal structure of the dynamics was preserved.

A larger difference appeared in how the final population state represented interval duration. With PSA, the projected distance grew approximately linearly with interval duration, whereas with MUA the relationship was nonlinear and saturated for longer intervals (Fig. 8D and Fig. S5I). IA fell in between: the projected distance still increased with duration, but less linearly than with PSA (Fig. S5I). The difference was also visible in the population trajectories, especially toward the end of int1 (Fig. S5K). We quantified it with ε95 and the relative RMS distance (see Methods) before and after Procrustes alignment (Fig. 8E); because this comparison focused on the population trajectories during the presentation of *int1* rather than on the whole trial, we used RMS distance instead of d². Unlike in the BDT, alignment reduced the discrepancy between PSA and MUA only modestly, from 14.3% to 11.5%. This fits the different relationships between interval duration and final population position, which no rotation or scaling of the trajectories can reconcile.

The overall shape of population dynamics can therefore be conserved across sorting methods while their geometry and task-related representations change. Even when population activity occupies relatively few significant dimensions, mixing neuronal responses can obscure how the circuit represents interval duration.

## Discussion

Spike sorting is often treated as a preprocessing step that can be skipped when population-level analyses are the goal. Our results show that whether it is needed depends on the inference being made. Applying nonlinear sorting (PSA), linear sorting (IA), or no sorting (MUA) to the same spikes, we found that sorting changed the number and firing rates of the identified units and their coding of stimulus periodicity and task variables. It also altered intrinsic timescales, trial-to-trial variability, neurometric sensitivity, the number of significant population dimensions, and the geometry of population dynamics. By contrast, the overall shape of the dynamics was largely conserved across representations. These effects were present in somatosensory and premotor cortex and in both tasks, but their magnitude depended on the area and on the type of information being quantified.

Multiunit threshold crossings support high-performance brain–computer interfaces (Gilja et al., 2015; Pandarinath et al., 2017; Willett et al., 2021, 2023) and preserve the low-dimensional trajectories that summarize population dynamics (Trautmann et al., 2019; Rossi-Pool and Romo, 2019). Our results agree with this view: after Procrustes alignment, the overall shape of PMv dynamics was nearly identical across representations. The same comparison, however, showed that the distances between population states, the number of significant dimensions, and the parametric representation of interval duration all depended on sorting. Decoding and manifold estimation rely on the dominant, shared structure of population activity, which pooling largely preserves. Analyses that ask which neurons carry information, how precisely they encode it, or how many independent dimensions a circuit uses depend instead on separating neurons with different response properties, which pooling undoes. Multiunit activity may therefore be sufficient when the goal is real-time decoding, whereas inferring how a circuit computes often requires single-neuron identity.

The difference between what pooling preserves and what it distorts can be traced to a single process: summing the spikes of neighboring neurons with heterogeneous response properties. When phase-locked neurons are pooled with neurons that lock at different phases or do not follow the stimulus at all, the periodic modulation of the mixed signal is diluted or partially cancels, and the mixed signal loses periodicity information while gaining rate information (Figs. 4 and S3). Pooling also changes the statistics of the signal. If neighboring neurons share positive noise correlations, the variance of their summed activity grows faster than its mean, which would raise the Fano factor of mixed activity; shared slow fluctuations would likewise lengthen its autocorrelation. This account fits the higher Fano factors and longer timescales we observed for IA and MUA. The same added variance propagates to population analyses: it raises the noise reference against which principal components are judged and lowers the number of significant dimensions. The structure that pooling removes is not necessarily noise, since correlated variability can enhance rather than impair the behavioral readout of population activity (Valente et al., 2021; Panzeri et al., 2022).

Because this process depends on how heterogeneous neighboring neurons are, its consequences should vary along the cortical hierarchy. In somatosensory cortex, periodicity coding in area 3b was largely robust to mixing, whereas area 1 lost much of the periodicity information carried by its single neurons. This difference matches the functional hierarchy between the two areas: compared with area 3b, area 1 has longer and more dispersed response latencies, greater trial-to-trial variability, and longer intrinsic timescales (Parra et al., 2022; Tauste Campo et al., 2024), all of which make neighboring neurons less likely to share the same temporal response. The same measures that define this hierarchy were themselves affected by sorting. In area 1, MUA lengthened the intrinsic timescale by about 40% relative to PSA, a bias comparable to the differences between adjacent areas that have been used to establish timescale hierarchies (Murray et al., 2014; Chaudhuri et al., 2015; Spitmaan et al., 2020; Rossi-Pool et al., 2021; Tauste Campo et al., 2024). Comparisons of timescales or variability across areas, or across studies, may therefore partly reflect differences in sorting rather than in circuit properties.

In PMv, where neighboring neurons show mixed selectivity and longer integration timescales, the consequences of mixing reached the population level. Mixing hid task-related dimensions in every dataset we examined: two of the five significant components in F5, the seventh component of the PMv population, which carried sustained delay activity, and in the TICT, the fourth component both during interval presentation and during the delay. Because these dimensions carried task-related dynamics, their loss changes the apparent complexity of the circuit. This loss relates to recent theoretical work showing that the ability to infer unrecorded neural activity from a recorded subset depends strongly on the effective dimensionality of the circuit. In connectome-constrained recurrent networks, Beiran and Litwin-Kumar (2025) showed that recordings from a subset of neurons can help constrain network dynamics and predict the activity of unrecorded neurons, but that the number of recorded neurons required for this inference depends on the dimensionality of the underlying activity. From this perspective, dimensionality also determines how many neurons must be recorded to capture the relevant degrees of freedom of a circuit, beyond describing population activity. If spike sorting changes dimensionality estimates, it may therefore alter our interpretation of how much of the circuit has been sampled and how much information remains in unobserved neurons.

In the TICT, mixing also changed how the population represented interval duration. With PSA, the final population state at the end of int1 moved approximately linearly with interval duration, consistent with the parametric code described for this circuit (Díaz et al., 2025). With MUA, this relationship saturated for longer intervals, and Procrustes alignment reduced the discrepancy between the two representations during *int1* only from 14.3% to 11.5%. The difference therefore cannot be explained by a rotation or rescaling of the trajectories; mixing altered the code itself. Other properties of the dynamics were preserved: trajectory speeds and the relative distances between trajectories followed similar patterns with all three representations. Mixed activity can thus reproduce the overall temporal structure of the population response while compressing the representation of longer durations. Studies that read elapsed or remembered time from population activity, and decoders designed to extract timing signals from multiunit recordings, may underestimate the resolution with which a circuit encodes long intervals.

The effects we report should be read as lower bounds on the cost of mixing. Our nonlinear sorting method was previously validated on synthetic datasets and on paired intracellular and extracellular recordings (Suárez-Barrera et al., 2025), and the units it isolates here are putative single neurons, some of which may still combine spikes from more than one cell. Any residual mixing in PSA would only reduce the differences between representations, so the true cost of pooling neurons is likely larger than we estimate.

Our results also point to two natural extensions of this work. First, our recordings were obtained with independently movable microelectrodes, in which each neuron is detected on a single channel. With high-density probes (Steinmetz et al., 2021), now also available for nonhuman primates (Trautmann et al., 2025), the same neuron can appear on several neighboring channels, so channel-wise multiunit signals count its spikes more than once and are no longer independent. Sorting that exploits the spatial footprint of each neuron across channels can assign those spikes to a single unit, so extending our nonlinear approach to multichannel waveforms is a direct next step. Second, we focused on measures derived from the spike counts and firing rates of individual units and populations. Measures defined between neurons, or between spikes and local field potentials, such as noise correlations, oscillatory phase, and spike–field coupling, depend explicitly on which neuron produced each spike and are likely to be at least as sensitive to mixing (Cohen & Kohn, 2011; Fries et al., 2001; Womelsdorf et al., 2007; Bastos et al., 2020). Quantifying how mixing affects these measures is the next step of this work.

These results suggest a simple rule for choosing between sorted and unsorted activity. Multiunit activity can support analyses that depend on the dominant, shared structure of population activity, such as real-time decoding or estimating the overall shape of population trajectories. Analyses that depend on how individual neurons differ often require spike sorting: the coding of fine stimulus features such as periodicity, estimates of variability and intrinsic timescales, neurometric sensitivity, the number of task-related dimensions, and parametric representations such as that of interval duration. Because all three representations can be obtained from the same recordings, the cost of this choice is computational rather than experimental, and comparing sorted and unsorted activity offers a direct test of whether a given conclusion depends on it. This distinction may become more relevant as brain–computer interfaces move beyond decoding movement toward decoding cognitive variables, such as decisions or elapsed time, whose representations depend on the heterogeneity of individual neurons rather than on the dominant population signal.

## Methods

### Datasets and ethics

All analyses were performed on previously published recordings from four rhesus monkeys (*Macaca mulatta*), which are publicly available on Zenodo (Zainos et al., 2022, 2024a, 2024b; Suárez-Barrera et al., 2025b; Encinas-Rodríguez et al., 2025). No new experiments were performed for this study. In the original experiments, animals were handled in accordance with the standards of the National Institutes of Health and the Society for Neuroscience, and all protocols were approved by the Institutional Animal Care and Use Committee of the Instituto de Fisiología Celular, Universidad Nacional Autónoma de México (Parra et al., 2022; Andrade-Ortega et al., 2025; Díaz et al., 2025).

### Spike detection and waveform extraction

Raw extracellular recordings were processed offline using custom Python scripts. Signals were sampled at 30 kHz and band-pass filtered between 300 and 3,000 Hz using a fourth-order Butterworth filter applied in forward and reverse directions to avoid phase shifts. Background noise was estimated independently for each trial and electrode as the median absolute deviation of the filtered signal divided by 0.6745 (Ǫuian Ǫuiroga et al., 2004). For each electrode, the median noise estimate across trials was used to define a fixed detection threshold for the entire recording session.

Candidate spikes were identified as negative-going threshold crossings exceeding four times the estimated background noise. A minimum separation of 1.5 ms between detected events was imposed to avoid multiple detections of the same waveform. Detected events were aligned to their local negative peaks, and 1.6-ms waveform segments were extracted, extending from 0.5 ms before to 1.1 ms after each peak. Events without a complete extraction window were excluded. Spike times, trial identifiers, and waveforms were then stored for spike sorting and subsequent analyses.

### Spike sorting workflow

Spike sorting followed the procedure of Suárez-Barrera et al. (2025) and consisted of three steps: preprocessing, feature extraction, and clustering.

#### Data preprocessing

To reduce noise, waveforms were smoothed with a third-order Savitzky-Golay filter with a 5-sample window, which preserves the essential features of each spike. Waveforms were then interpolated with a piecewise cubic Hermite interpolating polynomial (PCHIP), which yields a smooth interpolation and preserves their shape; each spike was aligned to its minimum and downsampled back to the original number of samples, keeping the alignment.

#### Feature extraction

We applied UMAP (McInnes et al., 2018) to the preprocessed waveforms for nonlinear dimensionality reduction, with two output dimensions (n_components = 2), a minimum distance of zero between embedded points (min_dist = 0), and 25 nearest neighbors (n_neighbors = 25). We chose the number of neighbors by comparing several values across different sets of sessions, keeping the minimum distance at zero to preserve the density of the data in the low-dimensional space. Dimensionality reduction was performed separately for each electrode, session, and recording area, in both the BDT and the TICT. UMAP draws on graph theory and differential geometry to capture both the local and the global arrangement of the data. Unlike methods such as t-SNE, UMAP preserves this global structure, which makes it well suited to large, high-dimensional datasets, so the low-dimensional embedding remains a faithful representation of the waveforms (McInnes et al., 2018).

We also applied principal component analysis (PCA) to the same waveforms. PCA defines a new coordinate system in which a few axes capture most of the variance: the 48 samples of each waveform are reduced to a few principal components (PCs), of which we always used the first two. Each PC explains the largest possible variance while remaining orthogonal to the previous ones. The PCs were derived from the covariance matrix estimated across all waveforms and time points:

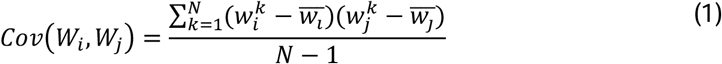

where *N* is the number of waveforms, 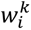 is the amplitude of the *k*-th waveform at time point *i*, and *w̄_j_* is the mean amplitude at time point j across all waveforms.

Diagonalizing this matrix (*Cov*(W*_i_*, W_j_)) yields the PCs (the columns of V) and the variance each one explains (the diagonal of Λ, a diagonal matrix with positive elements); PCs are ordered from highest to lowest variance. The projection of the *l*-th waveform onto the *k*-th PC is:

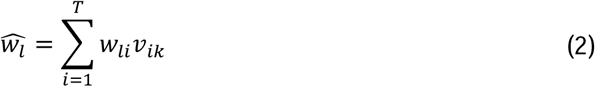

where *T* is the number of time points per waveform and *v_ik_* is the *i*-th element of the *k*-th PC; 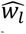 is the projection of waveform *w_l_* onto the *k*-th PC. Each PC is therefore a linear combination of the amplitudes at different time points of the waveform, and the contribution of each time point to a given (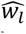) is given by the *i*-th element of the *k*-th column of the eigenvector matrix V. As shown in Suárez-Barrera et al. (2025), this and other linear methods capture the largest differences between waveforms but often merge the spikes of low-firing neurons with those of more active neurons, which dominate the variance.

#### Clustering

We clustered each embedding, obtained with UMAP or PCA, using HDBSCAN (Campello et al., 2013), with parameters adapted to the density of each embedding. HDBSCAN builds a cluster dendrogram from the connectivity between points, with the help of a minimum-distance parameter. By condensing the dendrogram, it identifies stable clusters, descending the hierarchy until it reaches an optimal density threshold and selecting the most prominent clusters. HDBSCAN handles noise natively: points that lie outside dense regions are labeled as noise. The parameters *min_cluster_size* and *min_samples* were adjusted linearly to the number of spikes recorded by each electrode in a session, within the ranges 40–70 and 90–175, respectively, for the embeddings obtained with UMAP, and within the ranges 100–200 and 40–80, respectively, for PCA. The *cluster_selection_epsilon* parameter was kept at 0 for UMAP, since the clusters were consistently compact and well-defined, while for PCA it was set to 0.2, since the generated clusters did not maintain a completely homogeneous density. Each cluster obtained in the embedding of each method was further validated by its interspike interval (ISI) violations and waveform stability: clusters with more than 3.5% of ISIs shorter than a 1.5-ms refractory period were excluded, as were spurious clusters whose activity never exceeded a firing rate of 2 Hz. Units obtained with UMAP were classified as putative single-unit activity (PSA), and those obtained with PCA as intermingled activity (IA), because they mix signals from low- and high-activity neurons. In addition, every analysis included the result of omitting spike sorting and only preprocessing the recorded signals, that is, multiunit activity (MUA), which comprises all spikes detected on each channel, including those left unassigned by HDBSCAN.

### Analysis of neuronal activity

#### Firing rate

Firing rates were computed across all correct trials of each neuron in each cognitive task (in the BDT, hits and correct rejections), using overlapping causal rectangular windows of 200 ms with 20-ms steps. For S1 neurons (areas 1 and 3b) in the BDT only, we used 50-ms windows with 10-ms steps across all trials. The firing rate of each neuron was averaged over the baseline period of each task, that is, before stimulation. To compare the distributions of mean firing rates, we built cumulative distribution functions for the units classified as PSA, IA, and MUA within each recording area.

#### Responsivity

To restrict the analyses to neurons that responded during the BDT and the TICT, we computed a z-score for each neuron (*n*) in each time window (*t*) relative to its baseline activity (*r_n_*⌃_,*basal*_):

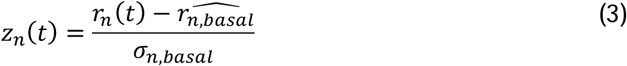

The baseline firing rate was obtained by averaging activity from −1 to 0 s. This window was applied to all trials, since every trial included at least 1 s of activity before stimulus onset.

S1 neurons (areas 1 and 3b) were classified as task-involved when their activity exceeded 2*σ_n_*_,*basal*_ in at least one time window during the stimulation period (0–0.5 s) for at least one suprathreshold stimulus. Task-involved S1 units that carried significant periodicity information (see *Mutual information*) were further classified as phase-locked.

For PMv neurons in the BDT, we used an analogous criterion: units were classified as task-involved when their activity exceeded 2*σ_n_*_,*basal*_or fell below −2*σ_n_*_,*basal*_for at least three consecutive time windows. This criterion had to be met for at least two suprathreshold amplitudes within the full task interval (0–3 s). In the TICT, it had to be met for at least two classes (distinct interval pairs) within the full task period.

#### Mutual information (MI)

For each neuron, we quantified the relationship between firing rate (*r*(*t*)) and the presented stimuli (*s*) in each time window using Shannon mutual information (Diaz-deLeon et al., 2022; Andrade-Ortega et al., 2025), a measure that can capture nonlinear dependencies between the two variables. To compute *I*(*r*; *s*), we used the firing-rate distributions conditioned on each stimulus condition (*P*(*r*|*s*)) together with the overall firing-rate distribution (*P*(*r*)), obtained by pooling trials across all conditions:

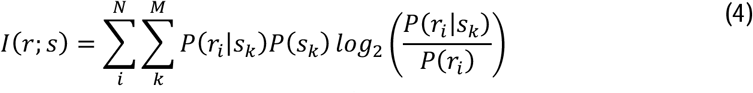

For the TICT, the information about interval duration (*I_int_*) was computed from the firing-rate distributions associated with each interval (*P*(*r*|*int*)) and the overall firing-rate distribution (*P*(*r*)), obtained by pooling trials across all interval values.

For S1 neurons in the BDT, tactile information (*I_tac_*) was computed from the firing-rate distributions associated with tactile stimuli (*P*(*r*|*tac*)) together with those obtained in the absence of a stimulus (*P*(*r*|*abs*)). The overall distribution (*P*(*r*)) was computed by pooling trials with and without tactile stimulation.

For PMv neurons in the BDT, modality information (*I_mod_*) was computed from the firing-rate distributions in stimulus-absent and tactile trials and, in addition, in acoustic trials (*P*(*r*|*acu*)). *P*(*r*) was obtained by pooling trials from all three conditions.

To obtain periodicity information (*I_period_*) in S1 neurons, firing rates were first estimated with a causal Gaussian kernel (*σ* = 25*ms*) to obtain a smooth time-resolved representation. A 200-ms sliding window was applied to the single-trial firing rate; within each window, the mean was subtracted, a Hamming window was applied, and a short-time Fourier transform was computed. The relative power in the 18–22 Hz band in each window was taken as its periodicity coefficient (*coef*). This yielded, for each tactile amplitude, a time-resolved distribution of periodicity coefficients (*P*(*coef*|*tac*)), from which we computed *I_period_* between 20-Hz periodicity and stimulus intensity:

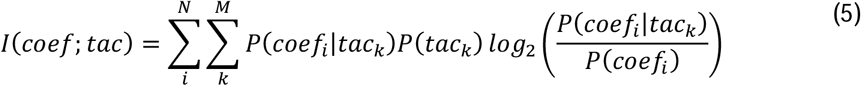

For each variable, significance was assessed for each neuron and time window. We performed 1,000 permutations of the trial labels and computed the information value for each permutation; the information in a given window was considered significant if the probability that a permutation yielded an equal or greater value was below 0.05 (P < 0.05). The number of neurons with significant mutual information was then corrected for multiple comparisons across neurons and time windows with a cluster-based method (Maris & Oostenveld, 2007), retaining only contiguous significant time windows whose size exceeded a defined threshold.

In all cases, mutual information values were also corrected for limited sampling. We permuted the class labels of the trials 1,000 times, computed time-resolved mutual information for each permutation, and estimated the average information expected from finite sampling:

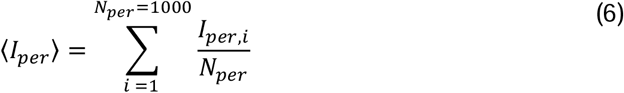

The corrected information was obtained by subtracting this average from the original value:

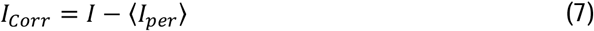

For the BDT, we compared the distributions of firing-rate and periodicity information, averaged over the stimulation period (0–0.5 s), within each S1 area (areas 1 and 3b) by building cumulative distribution functions for the units classified as PSA, IA, and MUA. Summed information (Figs. 4, 6, and S3) was obtained by adding the corrected information of the units with significant information at each time window.

#### Fano factor

For each neuron, we computed the variance and the mean of the spike count in each 50-ms time window of the baseline period (−1 to 0 s), and averaged their ratio across the windows of this period to obtain one mean Fano factor per neuron. To compare the distributions of baseline Fano factors across areas and methods, we built cumulative distribution functions for the units classified as PSA, IA, and MUA.

#### Coefficient of variation

We estimated the variability of the interspike intervals (ISIs) of each unit classified as PSA, IA, and MUA in the BDT, separately for each S1 area (areas 1 and 3b). For each trial, we took the activity in the period preceding possible stimulation (−0.5 to 0 s) and computed the distribution of ISI values. From these distributions, we computed the coefficient of variation of each unit as:

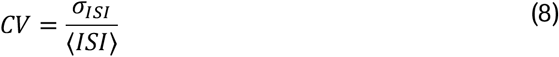

#### Autocorrelation

To characterize the autocorrelation of spike counts, we used the activity of the units classified as PSA, IA, and MUA in the prestimulus period (−0.6 to 0 s), separately for each S1 area (areas 3b and 1), following the procedure of Murray et al. (2014). The baseline period was divided into 40-ms time windows with 40-ms steps. For two time windows separated by a time lag Δ*t*, we computed the across-trial correlation of spike counts, *ρ*(Δ*t*). The resulting autocorrelation function was fitted by least squares with a biexponential decay function with an offset:

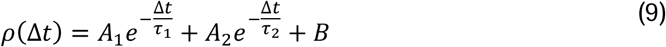

Following Spitmaan et al. (2020), who showed that cortical neurons express multiple timescales, the fit included two exponential components: a fast component (*τ*1), which we took as the intrinsic timescale, and a slow component (*τ*2), which absorbs slower fluctuations in the activity; *A*1 and *A*2 are their respective amplitudes. The offset (*B*) reflects the contribution of timescales much longer than our observation window (Murray et al., 2014). We performed this fit at both the population and the single-unit level. The error of the fitted parameter *τ*1 was estimated with a population bootstrap: synthetic datasets were created by resampling the neuronal population with replacement, so that each neuron could appear more than once or not at all in each resample. A new fit was computed for each synthetic dataset, yielding a distribution of *τ*1. The significance of the difference between methods was estimated from these bootstrap distributions, as the proportion of resamples in which *τ*1 for PSA was equal to or greater than that for MUA.

We compared the distributions of the fitted parameter *τ*1 across areas (areas 1 and 3b) by building cumulative distribution functions for the units classified as PSA, IA, and MUA.

#### Neurometric curves

To compute response distributions and neurometric detection curves, we used the units classified as PSA, IA, and MUA with significant periodicity coding. For each trial, we took the periodicity coefficient at 20 Hz during the stimulation period, computed as described above (relative power in the 18–22 Hz band; see Mutual information). Using stimulus-absent and tactile trials, neurometric curves were computed as the proportion of trials in which the maximum periodicity coefficient reached or exceeded a criterion (de Lafuente and Romo, 2005). For each neuron, this criterion was chosen to maximize the number of correct trials (hits and correct rejections). For each amplitude, the proportion of trials reaching or exceeding the criterion was averaged across neurons to obtain the population neurometric curve. From logistic fits, we computed the neurometric detection threshold of each method as the amplitude at which the probability of a “yes” response was 0.5 (Salinas et al., 2000).

#### Principal Component Analysis (PCA)

The main goal of PCA is to define a low-dimensional subspace that captures most of the variance of the high-dimensional neural state space. The number of significant dimensions drops markedly, from as many dimensions as neurons (*N*) to a few axes that concentrate most of the variance (Rossi-Pool & Romo, 2019).

To characterize how population activity covaries across classes over time, we performed PCA on F5 neurons (Fig. 7C) and on the whole PMv population (Figs. 8 and S4) during the BDT, using the threshold (1) and suprathreshold (2) classes of the two stimulus modalities (tactile and acoustic), together with the stimulus-absent condition (1 class). The covariance across classes was computed over time: from 0 to 3.5 s for the whole BDT, for the stimulation period (0 to 0.5 s), and for the delay (0.5 to 2.5 s). The resulting projection axes are mutually orthogonal unit vectors, ordered by the variance they explain: the first captures the most variance, the second the most of what remains, and so on. The projection axes were computed from the covariance matrix of the data:

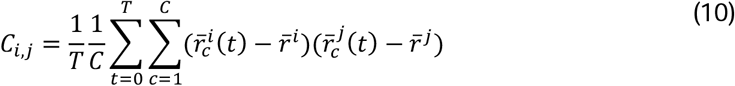

where *T* is the set of time windows of the interval analyzed, *C* is the number of classes (7 in this task), 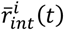 is the mean firing rate of neuron *i* in time window *t* and class *c*, and *r̄^l^* is the mean firing rate of neuron *i* across all time windows and classes.

In PMv during the TICT, to characterize how population activity varies across interval values (*int*) over time (*t*), we computed PCA for each method from the covariance matrix:

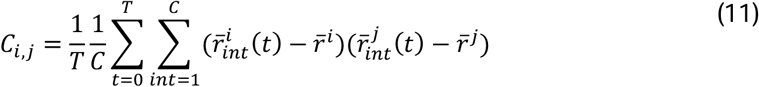

where *T* is the number of time windows in the period considered, *C* is the number of different intervals recorded in each experiment (7 in this task), *r̄^i^_int_* (*t*) is the mean firing rate of neuron *i* for interval int at time *t*, and *r̄^i^* is the mean firing rate of neuron *i* across all intervals and windows. We used neuronal activity from the onset of the first interval until 200 ms after its end; that is, only windows that included the interval presented on each trial were summed. These additional 200 ms are necessary because the response latency of PMv exceeds 150 ms (Rossi-Pool et al., 2020). We also analyzed the first delay period. In this case, the covariance matrix was computed from the time windows of this period on each trial, shifted forward by 200 ms to compensate for the response latency of PMv.

In both tasks (BDT and TICT), the projection of the N-dimensional data onto the k-th principal component (PC) is given by:

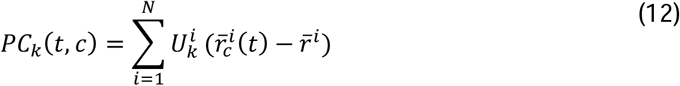

where *U^i^* is the *i*-th element of the *k*-th PC (*U_k_*) and *c* refers to each amplitude or interval (*int*) presented. PCs are therefore orthogonal linear combinations of the firing rates of individual neurons, uncorrelated with one another. The contribution of each neuron to *PC_k_* is thus given by *U^i^* .

#### Noise and dimensionality

To identify significant principal components in PMv in both tasks (BDT and TICT), we isolated the trial-to-trial fluctuations around task-related activity. For each neuron, the mean firing rate in each time window *t* across all trials of each class *c* or interval *int* was subtracted from the activity of each trial in that window and class. Using these residual fluctuations, we randomly selected 10 trials per neuron and computed the covariance matrix:

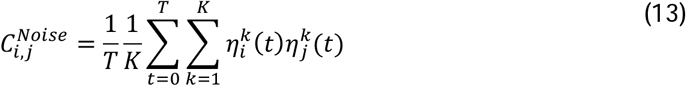

where *η^k^* is the residual firing rate at time *t* on trial *k* of neuron *i*. We applied PCA to this covariance matrix and obtained the variance captured by its first 10 principal components.

This procedure was repeated 100 times, and the variance captured by each of the first 10 components was averaged across repetitions. Principal components of the original covariance matrix whose variance exceeded that of the randomized trials were considered significant. We applied this procedure to different periods of the BDT (stimulation, 0–0.5 s; delay, 0.5–2.5 s; whole task, 0–3.5 s) and of the TICT (first interval and delay between intervals), to determine whether dimensionality, that is, the number of significant components, changed across periods with the different spike sorting strategies (PSA, IA, and MUA).

### Demixed Principal Component Analysis (dPCA)

The details of the algorithm and the mathematical justification of this method are given in Kobak et al. (2016). dPCA first separates neuronal activity according to predefined task variables to build marginalized covariance matrices, much like variable selection in a linear regression, and then applies standard PCA to these matrices. For the units classified in the BDT with each method (PSA, IA, and MUA) in PMv, we marginalized population activity (X) over the suprathreshold and threshold conditions (X*_t_*_,*c*_) of the tactile (3) and acoustic (3) modalities, excluding stimulus-absent trials. Leaving out stimulus-absent trials when computing the axes gives a clearer view of the population dynamics between the two modalities; activity from stimulus-absent trials can still be projected onto these axes (Andrade-Ortega et al., 2025).

To compute the demixed axes, the condition-independent component was subtracted during marginalization. The marginalized means are computed from the *N*-dimensional population activity:

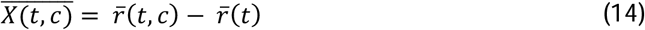

X̄_ϕ_ denotes the marginalized matrices, with *φ* ∈ {*t*, *c*} (the marginalization variable used in this study). Here, *r̄*(*t*) is the mean firing rate across all correct trials in each time window (pooling the 6 conditions, 3 tactile and 3 acoustic), whereas *r̄*(*t*, *c*) is the mean firing rate per condition (separating trials by the 3 conditions within each sensory modality, tactile or acoustic). After marginalization, dPCA finds decoding and encoding matrices for each X̄_*ϕ*_ by minimizing:

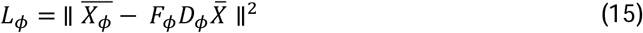

where X̅ is the centered population data matrix (that is, the mean activity of each neuron is 0). The minimization can be solved analytically by singular value decomposition. The components of *D_φ_* can be ordered by the variance they explain, as in standard PCA; the decoding axis that explains the most variance for variable *φ* is the first dPC. The significance of each dPC was assessed with the same noise-based criterion used for PCA (see Noise and dimensionality). For Fig. S4 and for the subsequent analysis of trajectory similarity in Fig. 8, we projected the *N*-dimensional data of a given condition (*r̄*(*t*, *c*), the mean firing rate per condition, computed by grouping trials by tactile amplitude, acoustic amplitude, or stimulus absence) onto the decoding axes (*k*) that explain the most variance for variable *φ*.

Fig. S4 shows the projections of neuronal activity onto the significant dPCs that explain the most variance for each condition, ordered by variance; these were used for the analyses in Figs. 8B and 8C. Modal axes are those that clearly separate tactile from acoustic stimuli, and we refer to them in Fig. S4 as modal dPCs; components that are independent of modality are called amodal dPCs.

#### Speed and Distance

In the TICT, for each spike sorting method (PSA, IA, and MUA), we computed the speed of and distances between neural trajectories using the Euclidean distance within the subspace spanned by the first significant PCs. For the absolute metrics, we compared trajectories directly at the same time points *t_i_*, measuring the differences between interval values (*int*) in the evolution of neuronal activity over time in the reduced space. From these differences we computed the speed shown in Fig. S5G.

We also computed the distance between the final positions of the population vector at the end of each interval (Fig. S5I). For this, we used the 1,000-ms interval as the reference. We first defined a projection axis from the difference vector between the reference interval and the 1,300-ms interval, and then computed the difference vector between the final position of the reference and the final positions of the other intervals.

For the relative metrics, we used kinematic analysis of neural trajectories (KiNeT), as described by Remington et al. (2018). KiNeT aligns neural trajectories according to their dynamics rather than strictly to external timing signals, which is especially important when comparing neural responses to stimuli of different durations or assessing how neural representations evolve during tasks that require temporal judgments. First, for each trajectory j, we found the time at which it was most similar to a reference trajectory, that is, the time *t*^j^ that minimized the Euclidean distance between the two trajectories in the PC subspace. Comparing this time of maximal similarity, *t*^j^, with the absolute time *t_i_* gave a measure of trajectory speed: a trajectory that reached the point of maximal similarity earlier than the reference was considered faster, and vice versa. This yielded the speed shown in Fig. S5H.

To compute the signed distance, we measured the Euclidean distance between trajectories at that relative time. We then assigned a sign to this distance based on the dot product between the difference vectors of the compared trajectories and the difference between the reference trajectory and the first trajectory. This captured both the magnitude of the differences between trajectories and their direction, that is, how they moved relative to one another (Fig. S5J).

#### Procrustes

To quantify the similarity between the population trajectories obtained with the different spike sorting methods (PSA, IA, and MUA), we performed a Procrustes analysis using the SciPy implementation. This procedure was applied to the PMv trajectories during the BDT, using seven PCA components and seven dPCA components, and during the TICT, using four principal components for the first interval. The trajectories of each representation were arranged in matrices X, *Y* ∈ ℝ^(*N*×*d*)^, where *N* is the number of population states, defined by experimental condition and time, and *d* is the number of components used. Correspondence between the states of the two representations was preserved in all comparisons.

To compute the discrepancy before alignment, each matrix was normalized by its Frobenius norm and the squared distance was computed:

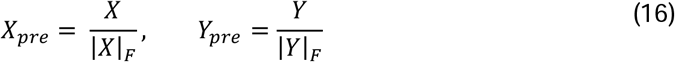

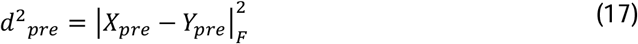

Procrustes alignment then centers both matrices by subtracting their centroids, normalizes them by their Frobenius norms, and finds the transformation that minimizes the sum of squared differences between corresponding states. The normalized matrices were defined as:

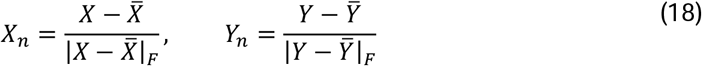

where X̅ and Y̅ are matrices whose elements are the centroids of X and *Y*. We determined an orthogonal matrix (*R*), which allows rotations and reflections, and a global scale factor (*s*) by minimizing:

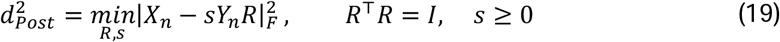

To express the aligned trajectories in the original coordinate system and units of the reference representation, the normalization and centering applied during Procrustes were inverted. The resulting transformation was:

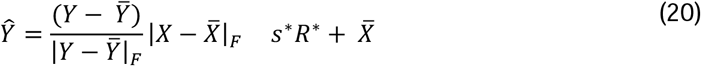

where (s*) and (*R\**) are the optimal scale factor and orthogonal transformation. This allowed the MUA trajectories to be represented in the PCA coordinate space of PSA while preserving the scale of the reference geometry.

We also computed 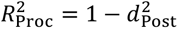 as a measure of geometric similarity after alignment.

We also expressed the square root of this discrepancy as the root-mean-square (RMS) error between the aligned trajectories, relative to the RMS size of the reference geometry.

***Geometric distortion (ε***_95_)

To assess how well the internal distances between population states are preserved across representations, we used a robust variant, *ε*_95_, of the geometric distortion ε proposed by Trautmann et al. (2019). Unlike Procrustes, this measure compares the distances between all pairs of states within each geometry, regardless of the orientation of its axes. For each pair of states (*i*, j), we computed the Euclidean distances *d_X_*(*i*, j) and *d*_F_(*i*, j) and their ratio:

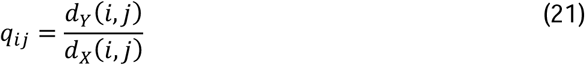

In its original form, the distortion is defined as the smallest maximum relative error that can be achieved with a single global scale factor (*λ*):

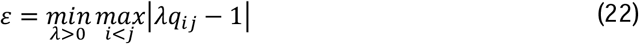

The optimal scale factor is obtained by balancing the errors associated with the smallest and largest ratios, *q*_min_ and *q*_max_:

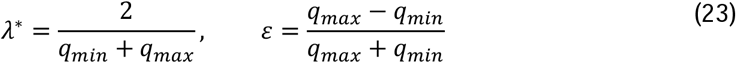

A value of *ε* = 0 indicates that all distances between states are preserved up to a common scale factor, and larger values indicate greater geometric distortion. Because *ε* depends on the most extreme ratios, it can be sensitive to small absolute differences between very close states, so a small fraction of pairs can fully determine its value. We therefore used *ε*95, the smallest maximum relative error that can be guaranteed for at least 95% of state pairs after optimizing a global scale factor. To compute it, the *M* ratios (*q_i_*_j_) were sorted in ascending order and a minimum number of pairs was set (*k* = ]0.95*M*]). We then evaluated every window of *k* consecutive ratios in the sorted distribution and selected the one with the smallest distortion:

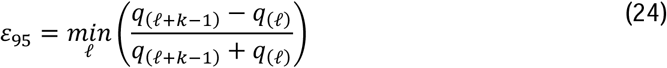

where *q*_(_*_P_*) is the ratio at position *P* of the sorted distribution. For the selected window, the optimal scale factor was computed from its minimum and maximum values. This procedure estimates the distortion that characterizes at least 95% of state pairs, keeping the global scale optimization while reducing the influence of extreme values. To avoid the influence of numerically null distances, pairs whose distance was practically zero in either representation were excluded, and trajectories were temporally subsampled at 100-ms intervals within each experimental condition, as a compromise between temporal resolution and the representation of temporally adjacent states.

#### Statistical analysis

To compare the cumulative distribution functions of mutual information, single-unit decay constants (*τ*), baseline Fano factor, and coefficient of variation (*CV*) across the three classification methods (PSA, IA, and MUA) within each area, each classified unit was treated as an independent observation. Differences between pairs of distributions were assessed with the Kolmogorov–Smirnov (*KS*) statistic, defined as the maximum absolute difference between the two cumulative distribution functions. Significance was estimated by bootstrap: the observations of the two groups were pooled, and 10,000 resamples with replacement were drawn with the same sizes as the original groups. The p value was computed as the proportion of resamples with a statistic equal to or greater than the original.

Confidence intervals and standard errors of neurometric thresholds were estimated with a population bootstrap. For each spike sorting method (PSA, IA, and MUA), starting from the neurometric curves of individual neurons, we drew 10,000 resamples with replacement, keeping the original number of neurons in each sample. For each sample, we computed the average population neurometric curve and fitted it with a sigmoid function. The neurometric threshold was defined as the stimulus amplitude at which the fitted detection probability was 0.5. This yielded a bootstrap distribution of neurometric thresholds for each method; the standard error was the standard deviation of this distribution, and 95% confidence intervals were obtained from its 2.5th and 97.5th percentiles.

## Use of large language models

A large language model (Claude, Anthropic) was used to assist with language editing. All content was reviewed, revised, and verified by the authors, who take full responsibility for the manuscript.

## Data availability

The biological results of the recordings analyzed here were previously described using publicly available datasets (Zainos et al., 2022, 2024a, 2024b). The complete electrophysiological signals were also used in the development of the UMAP-based sorting method and are publicly available on Zenodo (Suárez-Barrera et al., 2025; Suárez-Barrera et al., 2025b; Encinas-Rodríguez et al., 2025).

## Code availability

The code for spike sorting, firing rates, and mutual information is available at https://doi.org/10.5281/zenodo.17575203 (Suárez-Barrera et al., 2025). Scripts to classify neurons, compute firing rates, analyze coding, and perform PCA and dPCA are available with the bimodal detection task dataset (Zainos et al., 2024a) and at https://github.com/Rossi-Pool-Lab/Bimodal-Detection-analysis and https://github.com/BernardoAO/BDT-VPC. The code for the analyses introduced in this study will be deposited on GitHub (https://github.com/Rossi-Pool-Lab) upon publication.

## Acknowledgements

We thank Manuel Alvarez for technical assistance. This work was supported by grants from the Secretaría de Ciencia, Humanidades, Tecnología e Innovación (SECIHTI, CBF-2025-I-2054) and from the Dirección General de Asuntos del Personal Académico, UNAM (PAPIIT-IN203825), to R.R.-P.

## Author contributions

R.R.-P. conceived and designed the study. H.D., A.Z., L.B., N.E.-R., and D.S.-B. curated the data. L.B., N.E.-R., and D.S.-B. developed the software and analyzed the data. R.R.-P., L.B., N.E.-R., and D.S.-B. prepared the figures. R.R.-P., N.E.-R., and D.S.-B. wrote the manuscript. R.R.-P. supervised the project and acquired funding. All authors reviewed and approved the final manuscript.

---

**Figure S1.**
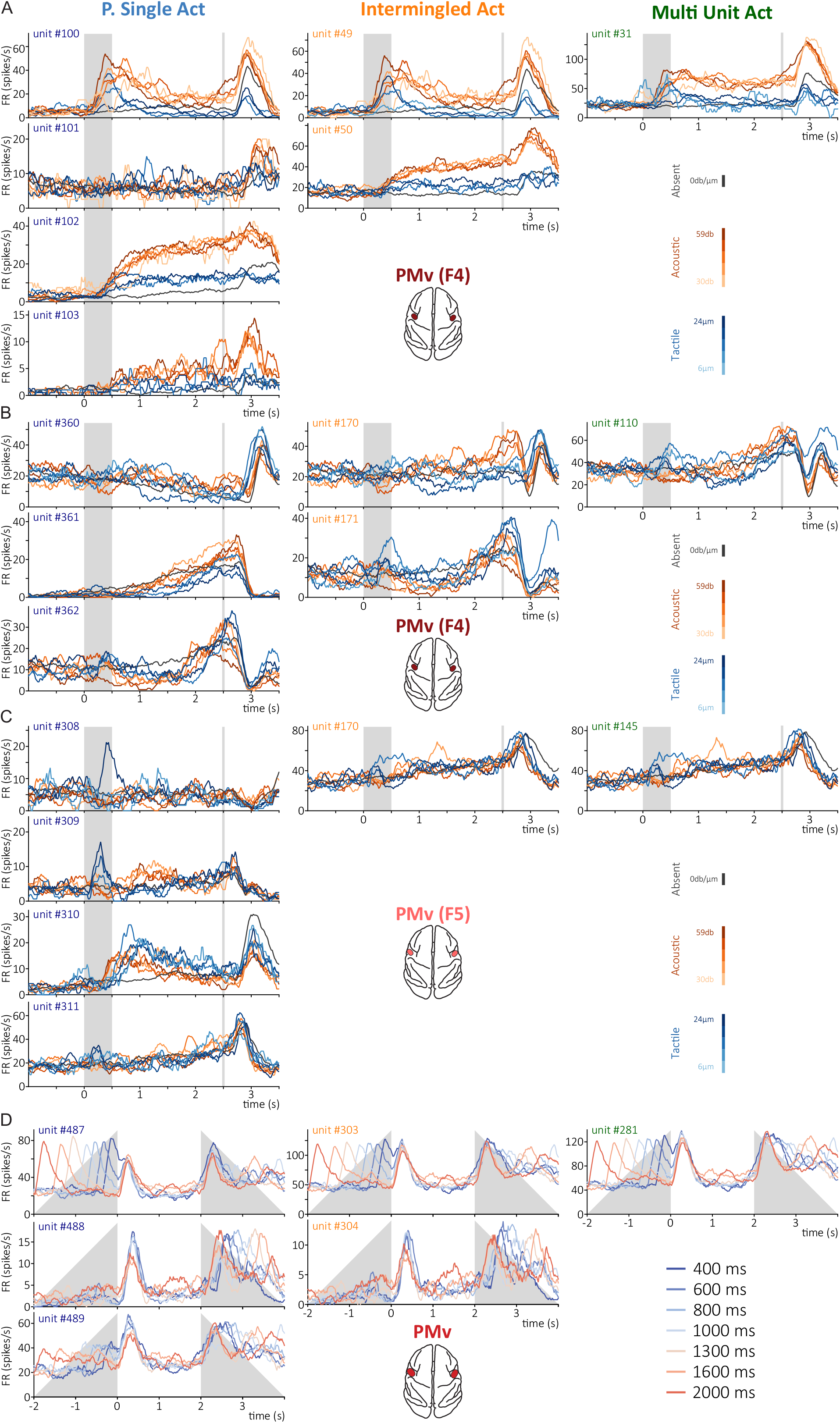
Examples of neuronal activity resolved by each spike sorting approach. (A–D) Firing rates of units identified with PSA (left), IA (middle), and MUA (right) in the same recordings. (A, B) PMv (F4) during the BDT. PSA identified four putative neurons in A and three in B, against two IA units and one MUA signal in each case. Neurons with different sensory preferences and temporal profiles contributed to the same mixed activity. (C) PMv (F5) during the BDT. PSA identified four putative neurons with distinct response profiles, whereas IA and MUA each yielded a single unit. In A–C, black traces are trials without stimulation, blue traces tactile stimuli (6 and 24 μm), and orange traces acoustic stimuli (30 and 59 dB). Gray shading marks the stimulus period; vertical gray lines mark the response cue. (D) PMv during the TICT. PSA identified three putative neurons, IA two units, and MUA one signal. Colors from blue to red indicate int1 duration (400–2,000 ms), and gray shading marks the variable timing of int1 and int2.

**Figure S2.**
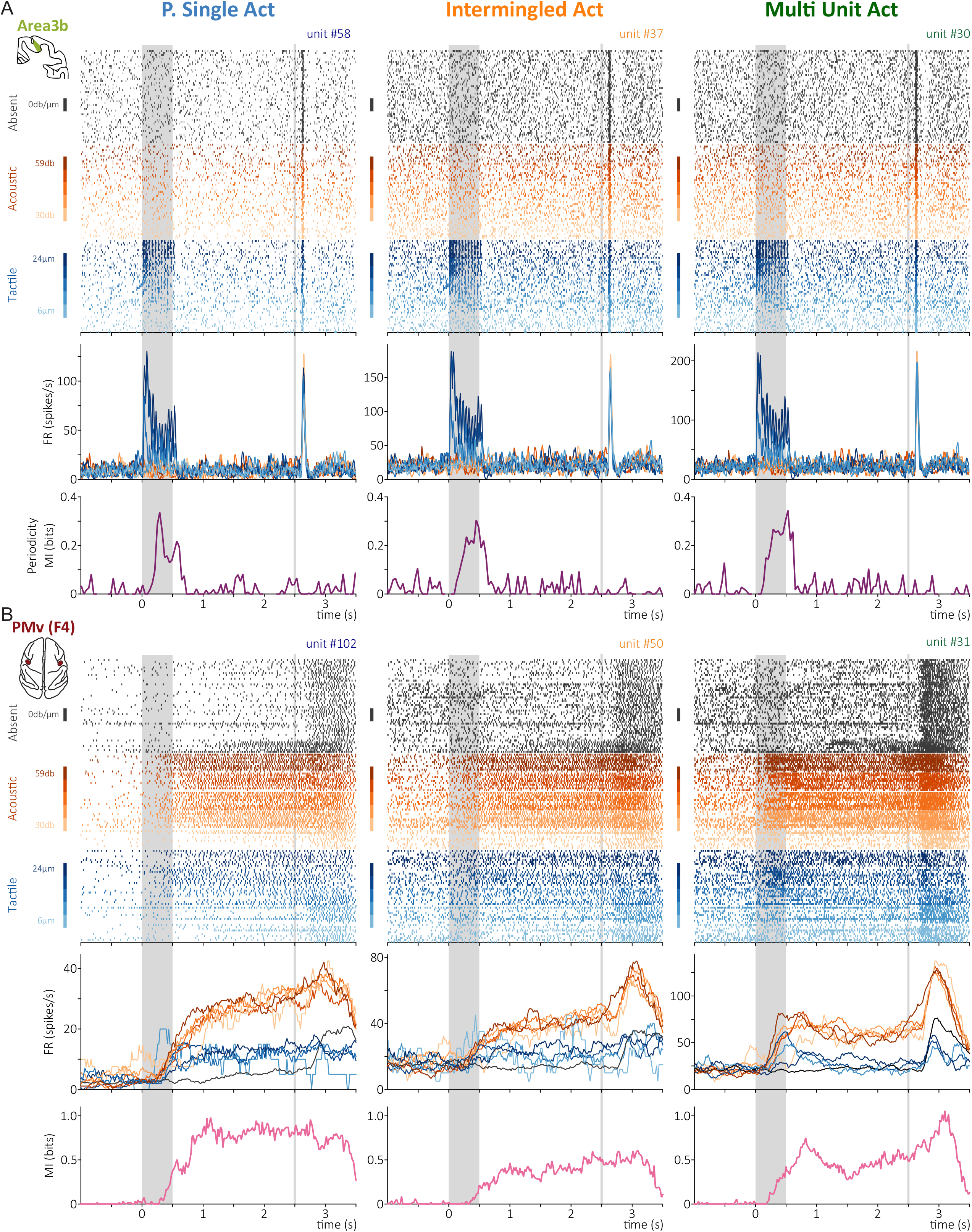
Additional examples of the effects of spike sorting on neuronal coding. (A) Area 3b during the BDT, organized as in Fig. 3A: spike rasters, firing rates, and MI about stimulus periodicity for PSA (left), IA (middle), and MUA (right). The three representations differed in firing rate and in the time course of periodicity information. (B) PMv (F4) during the BDT, with MI about stimulus modality in the bottom panels. The PSA neuron showed sustained activity and modality information, driven mainly by acoustic trials, throughout the delay. This information was reduced in IA, whereas MUA also included activity from neurons responding near the response period, producing a sharp rise in information at that time. Colors, shading, and time alignment as in Fig. 3.

**Figure S3.**
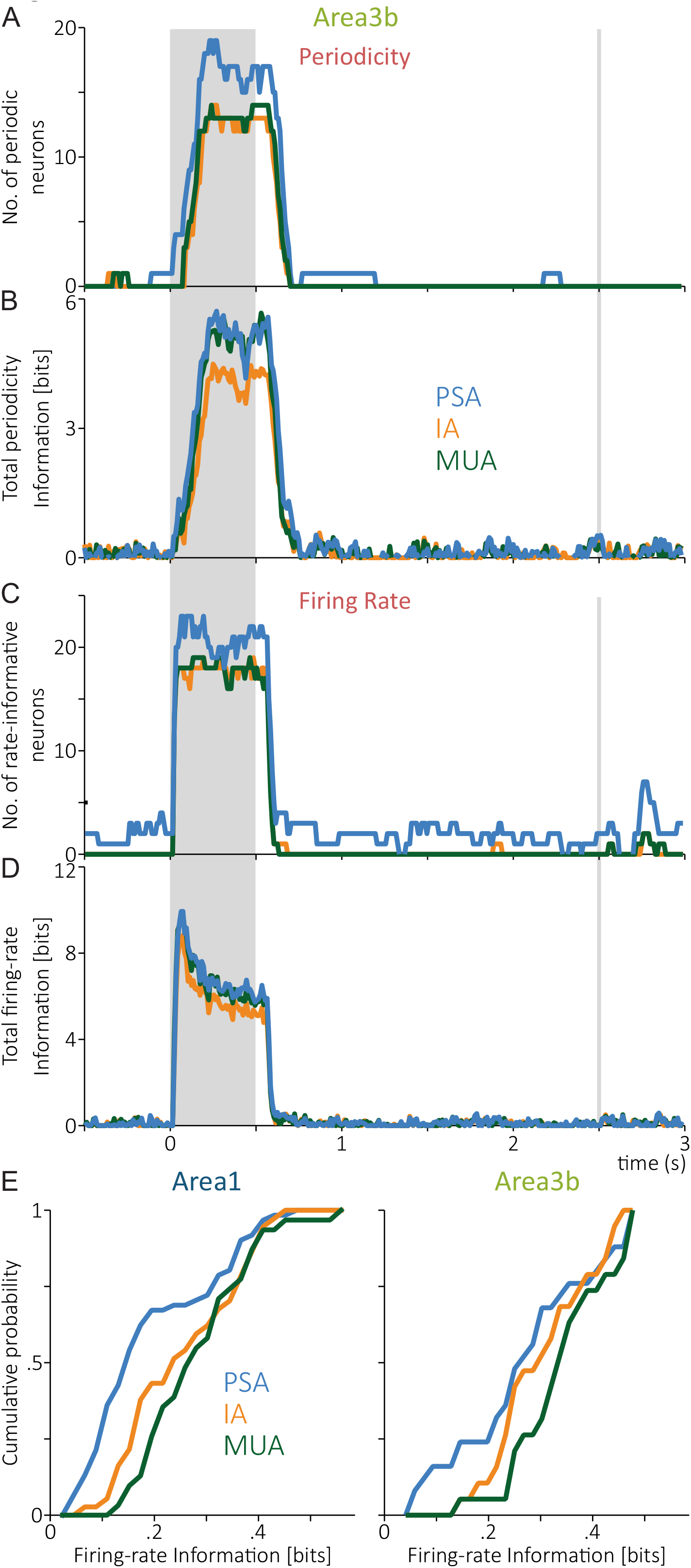
**Effects of spike sorting on periodicity and firing-rate coding**. (A–D) Area 3b during the BDT, for PSA (blue), IA (orange), and MUA (green), organized as in Fig. 4A–D. (A) Number of units with significant periodicity information over time. (B) Total periodicity information, summed across units. (C) Number of units with significant firing-rate information about stimulus amplitude. (D) Total firing-rate information, summed across units. (E) Cumulative distributions of firing-rate information per unit in area 1 (left) and area 3b (right). Shading, response cue, and time alignment as in Fig. 4.

**Figure S4.**
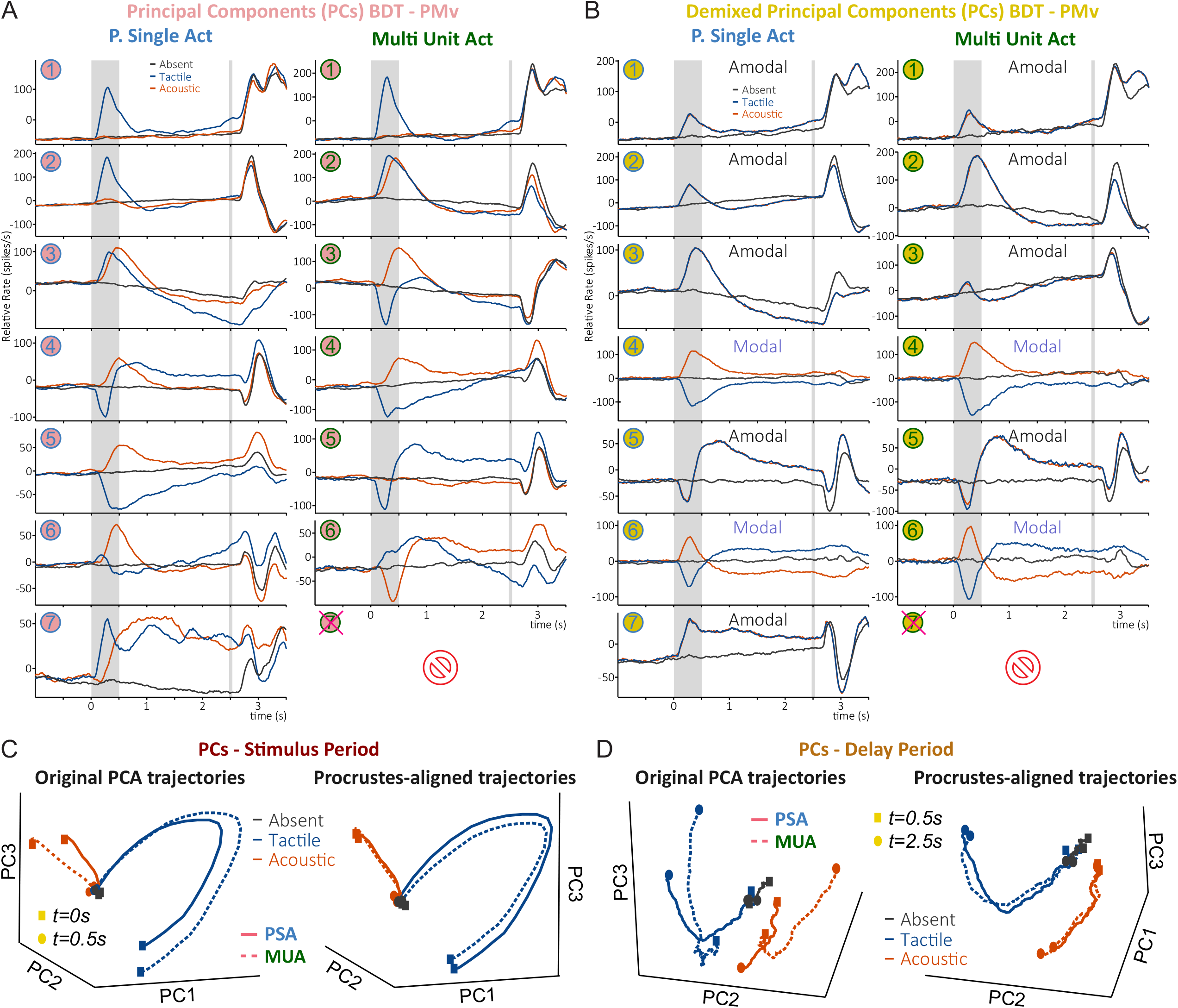
Effects of spike sorting on population dynamics during the BDT. (A) Time courses of the first seven principal components (PCs) of PMv activity for PSA (left) and MUA (right). Blue, tactile; orange, acoustic; black, stimulus absent. The crossed symbol marks the seventh MUA component, which was not significant. (B) The same analysis with dPCA. Components are labeled amodal when their activity is shared across stimulus modalities and modal when it depends on modality. The crossed symbol again marks the non-significant seventh MUA component. (C) Three-dimensional PCA trajectories during stimulus presentation (0–0.5 s), before (left) and after (right) Procrustes alignment. Solid lines, PSA; dashed lines, MUA. Squares and circles mark the start and end of the stimulus period. (D) The same comparison for the delay period (0.5–2.5 s). In A and B, time is relative to stimulus onset, gray shading marks the stimulus period, and vertical gray lines mark the response cue. All representations were computed from the same extracellular recordings.

**Figure S5.**
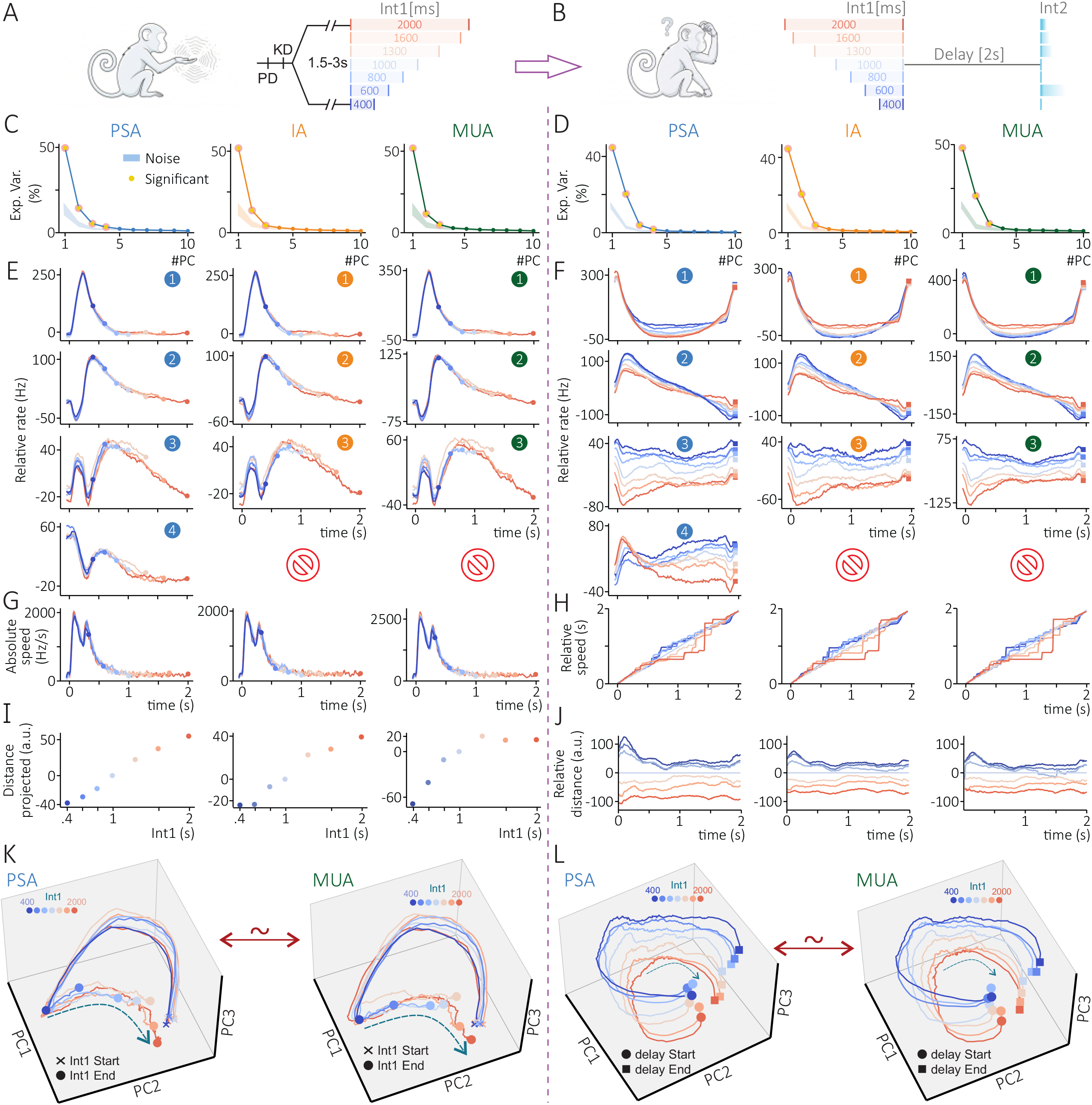
Effects of spike sorting on population dynamics during the TICT. (A, B) Task periods analyzed: representation of the first interval (int1; A) and maintenance of its duration during the following delay (B). Left column (C, E, G, I, K), int1; right column (D, F, H, J, L), delay. (C, D) Variance explained by the first ten principal components for PSA, IA, and MUA. Shaded regions show the variance expected from noise (see Methods), and yellow markers indicate significant components. (E, F) Time courses of the first four principal components. Colors indicate int1 duration, from 400 ms (blue) to 2,000 ms (red). The crossed symbol marks components that were not significant. (G) Absolute trajectory speed during int1. (H) Relative trajectory speed during the delay, estimated with KiNeT (Remington et al., 2018). (I) Projected distance between final population states as a function of int1 duration. (J) Relative distances between population trajectories during the delay, estimated with KiNeT. (K, L) Three-dimensional PCA trajectories for PSA and MUA during int1 (K) and the delay (L). In K, crosses and circles mark the start and end of int1; in L, circles and squares mark the start and end of the delay. Colors indicate int1 duration. All representations were computed from the same extracellular recordings.

## References

Andrade-Ortega, B., Díaz, H., Bayones, L., Alvarez, M., Zainos, A., Rivera-Yoshida, N., et al. (2025). Multistable bimodal perceptual coding within the ventral premotor cortex. Science Advances, 11(38), eadw5500.

Bastos, A. M., Lundqvist, M., Waite, A. S., Kopell, N., & Miller, E. K. (2020). Layer and rhythm specificity for predictive routing. Proceedings of the National Academy of Sciences, 117(49), 31459–31469.

Bayones, L., Zainos, A., Alvarez, M., Romo, R., Franci, A., & Rossi-Pool, R. (2024). Orthogonality of sensory and contextual categorical dynamics embedded in a continuum of responses from the second somatosensory cortex. Proceedings of the National Academy of Sciences, 121(29), e2316765121.

Beiran, M., & Litwin-Kumar, A. (2025). Prediction of neural activity in connectome-constrained recurrent networks. Nature Neuroscience, 28(12), 2561–2574.

Britten, K. H., Shadlen, M. N., Newsome, W. T., & Movshon, J. A. (1992). The analysis of visual motion: A comparison of neuronal and psychophysical performance. Journal of Neuroscience, 12(12), 4745–4765.

Buffalo, E. A., Fries, P., Landman, R., Buschman, T. J., C Desimone, R. (2011). Laminar differences in gamma and alpha coherence in the ventral stream. Proceedings of the National Academy of Sciences, 108(27), 11262–11267.

Campello, R. J. G. B., Moulavi, D., & Sander, J. (2013). Density-based clustering based on hierarchical density estimates. In J. Pei, V. S. Tseng, L. Cao, H. Motoda, & G. Xu (Eds.), Advances in Knowledge Discovery and Data Mining. PAKDD 2013. Lecture Notes in Computer Science (Vol. 7819). Springer.

Chaudhuri, R., Knoblauch, K., Gariel, M.-A., Kennedy, H., & Wang, X.-J. (2015). A large-scale circuit mechanism for hierarchical dynamical processing in the primate cortex. Neuron, 88(2), 419–431.

Churchland, M. M., Yu, B. M., Cunningham, J. P., Sugrue, L. P., Cohen, M. R., Corrado, G. S., et al. (2010). Stimulus onset quenches neural variability: A widespread cortical phenomenon. Nature Neuroscience, 13(3), 369–378.

Cohen, M. R., & Kohn, A. (2011). Measuring and interpreting neuronal correlations. Nature Neuroscience, 14, 811–819.

Cunningham, J. P., & Yu, B. M. (2014). Dimensionality reduction for large-scale neural recordings. Nature Neuroscience, 17, 1500–1509.

de Lafuente, V., & Romo, R. (2005). Neuronal correlates of subjective sensory experience. Nature Neuroscience, 8(12), 1698–1703.

Díaz, H., Bayones, L., Alvarez, M., Andrade-Ortega, B., Valero, S., Zainos, A., et al. (2025). Contextual neural dynamics during time perception in the primate ventral premotor cortex. Proceedings of the National Academy of Sciences, 122(6), e2420356122.

Diaz-deLeon, G., Alvarez, M., Bayones, L., Zainos, A., Zizumbo, J., Parra, S., et al. (2022). An abstract categorical decision code in dorsal premotor cortex. Proceedings of the National Academy of Sciences, 119, e2214562119.

Ecker, A. S., Berens, P., Keliris, G. A., Bethge, M., Logothetis, N. K., & Tolias, A. S. (2014). State dependence of noise correlations in macaque primary visual cortex. Neuron, 82(1), 235–248.

Elsayed, G. F., & Cunningham, J. P. (2017). Structure in neural population recordings: An expected byproduct of simpler phenomena? Nature Neuroscience, 20(9), 1310–1318.

Encinas-Rodríguez, N., et al. (2025). *Foreperiod neural activity recordings from ventral premotor (VPC), dorsal premotor (DPC), and secondary somatosensory (S2) cortices* [Data set]. Zenodo. 10.5281/zenodo.17569100

Felleman, D. J., & Van Essen, D. C. (1991). Distributed hierarchical processing in the primate cerebral cortex. Cerebral Cortex, 1(1), 1–47.

Fries, P., Reynolds, J. H., Rorie, A. E., & Desimone, R. (2001). Modulation of oscillatory neuronal synchronization by selective visual attention. Science, 291(5508), 1560–1563.

Gao, P., & Ganguli, S. (2015). On simplicity and complexity in the brave new world of large-scale neuroscience. Current Opinion in Neurobiology, 32, 148–155.

Gilja, V., Pandarinath, C., Blabe, C. H., Nuyujukian, P., Simeral, J. D., Sarma, A. A., et al. (2015). Clinical translation of a high-performance neural prosthesis. Nature Medicine, 21(10), 1142–1145.

Hernández, A., Zainos, A., & Romo, R. (2000). Neuronal correlates of sensory discrimination in the somatosensory cortex. Proceedings of the National Academy of Sciences, 97(11), 6191–6196.

Kobak, D., Brendel, W., Constantinidis, C., Feierstein, C. E., Kepecs, A., Mainen, Z. F., et al. (2016). Demixed principal component analysis of neural population data. eLife, 5, e10989.

Kohn, A., Coen-Cagli, R., Kanitscheider, I., & Pouget, A. (2016). Correlations and neuronal population information. Annual Review of Neuroscience, 39, 237–256.

Maris, E., & Oostenveld, R. (2007). Nonparametric statistical testing of EEG- and MEG-data. Journal of Neuroscience Methods, 164(1), 177–190.

Matelli, M., Luppino, G., & Rizzolatti, G. (1985). Patterns of cytochrome oxidase activity in the frontal agranular cortex of the macaque monkey. Behavioural Brain Research, 18(2), 125–136.

McInnes, L., Healy, J., Saul, N., & Großberger, L. (2018). UMAP: Uniform manifold approximation and projection. Journal of Open Source Software, 3(29), 861.

Murray, J. D., Bernacchia, A., Freedman, D. J., Romo, R., Wallis, J. D., Cai, X.-J., Padoa-Schioppa, C., Pasternak, T., Seo, H., Lee, D., & Wang, X.-J. (2014). A hierarchy of intrinsic timescales across primate cortex. Nature Neuroscience, 17, 1661–1663.

Newsome, W. T., Britten, K. H., & Movshon, J. A. (1989). Neuronal correlates of a perceptual decision. Nature, 341, 52–54.

Pandarinath, C., Nuyujukian, P., Blabe, C. H., Sorice, B. L., Saab, J., Willett, F. R., Hochberg, L. R., Shenoy, K. V., & Henderson, J. M. (2017). High performance communication by people with paralysis using an intracortical brain-computer interface. eLife, 6, e18554.

Panzeri, S., Moroni, M., Safaai, H., & Harvey, C. D. (2022). The structures and functions of correlations in neural population codes. Nature Reviews Neuroscience, 23(9), 551–567.

Papale, P., Wang, F., Self, M. W., & Roelfsema, P. R. (2025). An extensive dataset of spiking activity to reveal the syntax of the ventral stream. Neuron, 113(4), 539–553.e5.

Parra, S., Díaz, H., Zainos, A., Alvarez, M., Zizumbo, J., Rivera-Yoshida, N., et al. (2022). Hierarchical unimodal processing within the primary somatosensory cortex during a bimodal detection task. Proceedings of the National Academy of Sciences, 119(52), e2213847119.

Quian Quiroga, R., Nadasdy, Z., & Ben-Shaul, Y. (2004). Unsupervised spike detection and sorting with wavelets and superparamagnetic clustering. Neural Computation, 16(8), 1661–1687.

Remington, E. D., Narain, D., Hosseini, E. A., & Jazayeri, M. (2018). Flexible sensorimotor computations through rapid reconfiguration of cortical dynamics. Neuron, 98(5), 1005–1019.e5.

Romo, R., & Salinas, E. (2003). Flutter discrimination: Neural codes, perception, memory and decision making. Nature Reviews Neuroscience, 4(3), 203–218.

Rossi-Pool, R., & Romo, R. (2019). Low dimensionality, high robustness in neural population dynamics. Neuron, 103(2), 177–179.

Rossi-Pool, R., Vergara, J., & Romo, R. (2020). Constructing perceptual decision-making across cortex. In D. Poeppel, G. R. Mangun, & M. S. Gazzaniga (Eds.), The cognitive neurosciences (6th ed., pp. 413–427). MIT Press.

Rossi-Pool, R., Zainos, A., Alvarez, M., Zizumbo, J., Vergara, J., & Romo, R. (2017). Decoding a decision process in the neuronal population of dorsal premotor cortex. Neuron, 96, 1432–1446.

Rossi-Pool, R., Zainos, A., Alvarez, M., Parra, S., Zizumbo, J., Romo, R., et al. (2021). Invariant timescale hierarchy across the cortical somatosensory network. Proceedings of the National Academy of Sciences, 118(3), e2021843118.

Salinas, E., Hernández, A., Zainos, A., & Romo, R. (2000). Periodicity and firing rate as candidate neural codes for the frequency of vibrotactile stimuli. Journal of Neuroscience, 20(14), 5503–5515.

Spitmaan, M., Seo, H., Lee, D., & Soltani, A. (2020). Multiple timescales of neural dynamics and integration of task-relevant signals across cortex. Proceedings of the National Academy of Sciences, 117(36), 22522–22531.

Steinmetz, N. A., Aydin, C., Lebedeva, A., Okun, M., Pachitariu, M., Bauza, M., et al. (2021). Neuropixels 2.0: A miniaturized high-density probe for stable, long-term brain recordings. Science, 372, eabf4588.

Suárez-Barrera, D., Bayones, L., Encinas-Rodríguez, N., Parra, S., Monroy, V., Pujalte, S., et al. (2025). Efficient and reliable spike sorting from neural recordings with UMAP-based unsupervised nonlinear dimensionality reduction. PLOS Biology, 23(11), e3003527.

Suárez-Barrera, D., et al. (2025b). Electrophysiological recordings of neurons in S2, VPC, and DPC during cognitive tasks [Data set]. Zenodo. 10.5281/zenodo.17546569

Tauste Campo, A., Zainos, A., Vázquez, Y., Adell Segarra, R., Álvarez, M., Deco, G., et al. (2024). Thalamocortical interactions shape hierarchical neural variability during stimulus perception. iScience, 27, 110065.

Trautmann, E. M., Stavisky, S. D., Lahiri, S., Ames, K. C., Kaufman, M. T., O’Shea, D. J., Vyas, S., Sun, X., Ryu, S. I., Ganguli, S., & Shenoy, K. V. (2019). Accurate estimation of neural population dynamics without spike sorting. Neuron, 103(2), 292–308.e4.

Trautmann, E. M., Hesse, J. K., Stine, G. M., Xia, R., Zhu, S., O’Shea, D. J., et al. (2025). Large-scale high-density brain-wide neural recording in nonhuman primates. Nature Neuroscience, 28(7), 1562–1575.

Valente, M., Pica, G., Bondanelli, G., Moroni, M., Runyan, C. A., Morcos, A. S., Harvey, C. D., & Panzeri, S. (2021). Correlations enhance the behavioral readout of neural population activity in association cortex. Nature Neuroscience, 24(7), 975–986.

Vyas, S., Golub, M. D., Sussillo, D., & Shenoy, K. V. (2020). Computation through neural population dynamics. Annual Review of Neuroscience, 43, 249–275.

Willett, F. R., Avansino, D. T., Hochberg, L. R., Henderson, J. M., & Shenoy, K. V. (2021). High-performance brain-to-text communication via handwriting. Nature, 593, 249–254.

Willett, F. R., Kunz, E. M., Fan, C., Avansino, D. T., Wilson, G. H., Choi, E. Y., Kamdar, F., Glasser, M. F., Hochberg, L. R., Druckmann, S., Shenoy, K. V., & Henderson, J. M. (2023). A high-performance speech neuroprosthesis. Nature, 620, 1031–1036.

Womelsdorf, T., Schoffelen, J.-M., Oostenveld, R., Singer, W., Desimone, R., Engel, A. K., & Fries, P. (2007). Modulation of neuronal interactions through neuronal synchronization. Science, 316(5831), 1609–1612.

Zainos, A., Díaz, H., Rivera-Yoshida, N., Alvarez, M., Parra, S., Zizumbo, J., et al. (2022). Bimodal detection task dataset [Data set]. Zenodo. 10.5281/zenodo.7293495

Zainos, A., Díaz, H., Rivera-Yoshida, N., Alvarez, M., Andrade-Ortega, B., Bayones, L., Romo, R., & Rossi-Pool, R. (2024a). Bimodal detection task VPC [Data set]. Zenodo. 10.5281/zenodo.14361832

Zainos, A., Díaz, H., Alvarez, M., Bayones, L., Andrade-Ortega, B., Valero, S., & Rossi-Pool, R. (2024b). *Time perception VPC dataset* [Data set]. Zenodo. 10.5281/zenodo.14441503

